# Giant viruses encode vitamin K–based redox modules for lipid modification

**DOI:** 10.64898/2026.03.26.714578

**Authors:** Rebecca Collins, Julien Adréani, Dariana Chavez, Douglas B. Rusch, Dana Boyd, Philippe Colson, Bernard La Scola, Cristina Landeta

## Abstract

Viruses with large DNA genomes often carry auxiliary metabolic genes that reprogram host physiology, yet their contributions to host redox and membrane homeostasis remains poorly understood. Here we report the discovery and functional reconstitution of viral homologs of vitamin K epoxide reductase (VKOR) encoded by giant viruses. Using phylogenetic and genomic context analysis, we find that viral VKOR genes are frequently located adjacent to γ-carboxylase-like epoxidase and fatty acid desaturase domains, consistent with a putative modular redox pathway for membrane lipid modification. To investigate their function, we expressed viral VKORs in an *Escherichia coli* strain lacking disulfide bond–forming enzymes and examined both their membrane topology and activity. Remarkably, a minimal set of residue substitutions enabled proper membrane insertion and restored bacterial motility, demonstrating that viral VKORs are catalytically competent electron shuttles. Structural modeling supports their integration into the endoplasmic reticulum–like environment in the host. Finally, we show that VKORs and γ-carboxylase-like epoxidase-desaturases from *Fadolivirus* and *Yasminevirus* giant viruses are expressed during infection of *Vermamoeba vermiformis*, where they may couple vitamin K epoxidation to desaturation-driven lipid remodeling. These findings expand the known functional repertoire of giant viruses and uncover a previously unrecognized viral strategy for manipulating host redox metabolism and membrane composition.

**Significance statement:** Viruses with large DNA genomes often encode metabolic enzymes that reshape host physiology, yet their ability to control host redox and membrane composition has remained unclear. We discovered that giant viruses encode a vitamin K epoxide reductase and associated enzymes that together form a putative redox module for lipid remodeling. We reconstituted viral VKOR activity in *E. coli*, demonstrating that these enzymes are catalytically active. We also detected their expression, at the RNA and protein levels, during amoebal infection by two giant viruses. Our findings reveal a previously unrecognized viral strategy for coupling vitamin K redox cycling to fatty acid desaturation. This work broadens our understanding of how giant viruses manipulate host redox homeostasis and membrane architecture during infection.

## Introduction

Disulfide bonds are posttranslational modifications formed when electrons are removed from two cysteines, creating a stabilizing covalent linkage (1). These bonds help proteins fold and remain stable in oxidizing environments and are common in exported proteins across the bacterial cell envelope, endoplasmic reticulum (ER), Golgi, chloroplasts, and mitochondria (2). Cytoplasmic proteins in both prokaryotes and eukaryotes are generally reduced, with exceptions such as transient stress-induced disulfides (3, 4). Besides certain viruses, such as *Orthopoxvirus vaccinia* virus, encode enzymes that produce disulfide-bonded viral assembly proteins in the host cytoplasm (5, 6). Hyperthermophilic organisms may also contain cytosolic disulfide bonds, likely to stabilize proteins at high temperatures (7–9).

Disulfide bond formation proceeds through three redox steps involving catalytic cysteines. A disulfide-bond catalyst first oxidizes the client protein (Fig. 1a; e.g., PDI in eukaryotes, DsbA in bacteria) (10, 11). A membrane enzyme with its cofactor then regenerates the catalyst (Fig. 1a; e.g., Ero1/FAD in eukaryotes; DsbB/quinone in bacteria) (12, 13). Finally, the cofactor transfers electrons to other cellular processes, such as the electron transport chain in bacteria (Fig. 1a) (14, 15).

**Figure 1.**
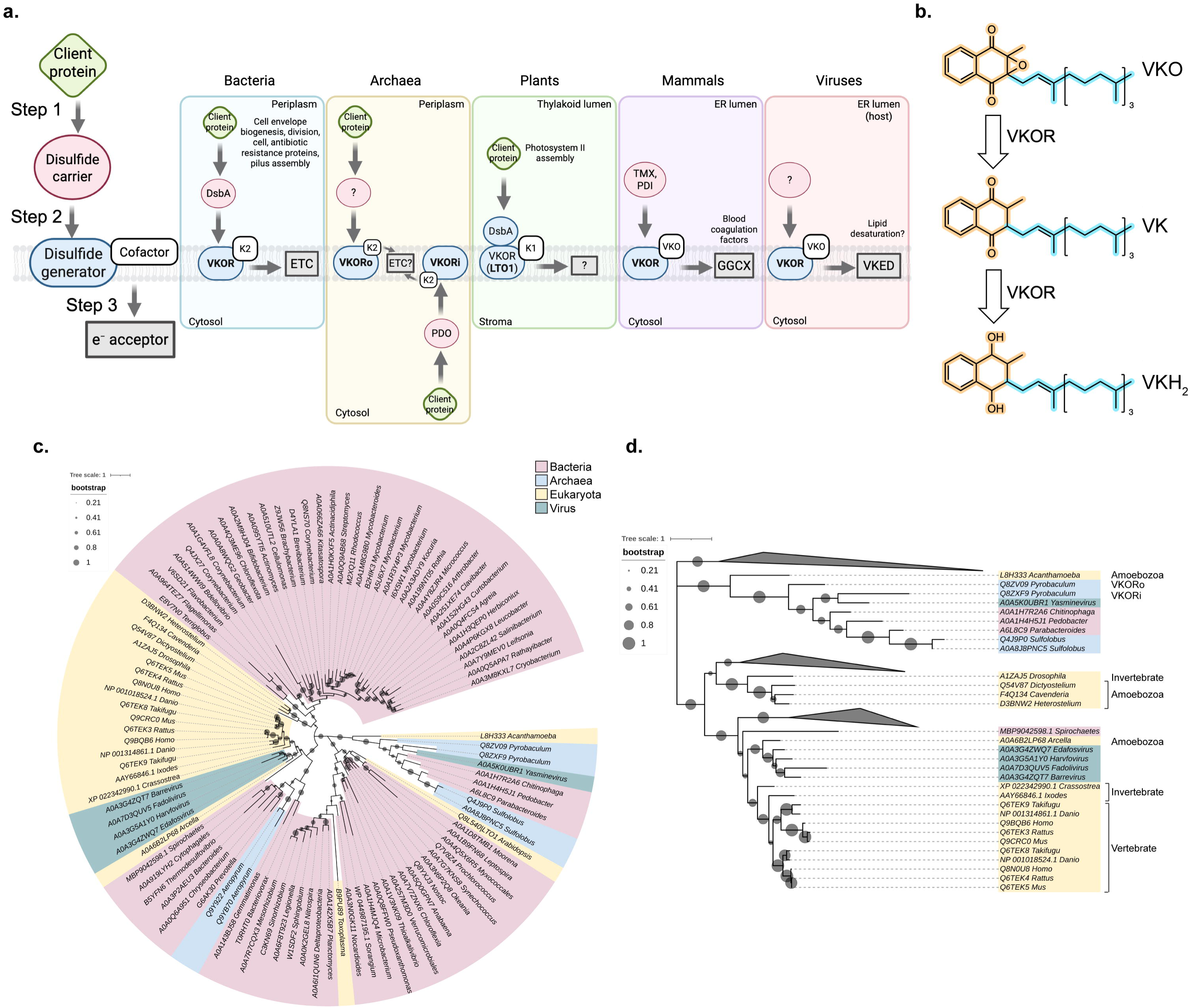
Giant viruses harbor VKOR enzymes. **a**. Redox steps of disulfide bond formation are conserved across all domains of life. See text for further details. e^-^, electron; K2, menaquinone or vitamin K_2_; K1, phylloquinone or vitamin K_1_; VKO, vitamin K epoxide; ?, unknown. **b**. Human VKORc1 can reduce vitamin K epoxide to vitamin K hydroquinone in two steps that each involves a two-electron transfer. **c**. Four viral VKOR proteins group together with VKOR from the amoeba *Arcella intermedia* but not with other amoebae. VKOR phylogenetic tree was done using protein sequences from 100 non-redundant species by a multiple sequence alignment using Muscle and MEGA12 (80). The phylogenetic reconstructions were inferred using the Maximum Likelihood method and Whelan-Goldman WAG model (+Freq) (81). The Newick tree was then visualized using iTol (https://itol.embl.de/). Uniprot identifiers are shown next to genera. **d**. A smaller subset of the tree was used for representatives of each family. Uniprot identifiers are shown next to the organism’s names. VKORo: facing outward VKOR; and VKORi: facing inward VKOR.

Vitamin K epoxide reductase (VKOR) is a conserved membrane enzyme in the second redox step, present across all domains except fungi (16, 17). Bacterial VKOR was identified in species where the enzyme is fused to DsbA (18, 19), such as *Synechococcus* sp. VKOR that can catalyze both the first and second steps (19, 20). Bacterial VKOR maintains DsbA oxidized by transferring the electrons to menaquinone (vitamin K_2_) (18, 21, 22). Bacterial VKOR participates in the folding of proteins required for cell envelope biogenesis, cell division and pilus assembly (Fig. 1a) (23, 24). Archaeal VKORs can face either membrane side, potentially enabling disulfide formation in both compartments (Fig. 1a) (25, 26). In plants, VKOR is also fused to a DsbA-like protein (named LTO1) and uses phylloquinone (vitamin K_1_) to support Photosystem II assembly in the thylakoid lumen of the chloroplast (Fig. 1a) (27–29). Vertebrate VKORc1, located in the ER, interacts with luminal oxidoreductases (TMX, TMX4, Erp18 and PDI) (30) but is only a minor contributor of disulfide bond formation (31). Instead, it drives vitamin K–dependent pathways such as blood coagulation by regenerating vitamin K hydroquinone (VKH_2_) for γ-carboxylation of clotting factors (Fig. 1a) (32, 33). To regenerate VKH_2_, VKORc1 reduces the vitamin K epoxide (VKO) first to quinone (VK), and then to hydroquinone (VKH_2_), with each reduction step involving two electrons (Fig. 1b) (34). Thus, one key difference between bacterial, archaeal and plantae VKOR orthologs with vertebrate VKOR is that the former proteins only enable the reduction of the quinone to the hydroquinone, since the epoxide forms are not known to be produced in these compartments (27, 35, 36). However, mycobacterial VKOR can catalyze the reduction of VKO and VK when expressed in mammalian cell lines (37). Thus, bacterial VKOR can respond to the cofactors present in the cellular environment. Vertebrates additionally encode VKORc1L1, essential for extrahepatic vitamin K metabolism and bone homeostasis (38). Phylogenetic analyses revealed five VKOR clades spanning bacteria, archaea, vertebrates, non-vertebrate eukaryotes, and photosynthetic organisms (17).

The discovery of more complex and diverse giant viruses in recent years has deepened our perception of the virosphere since the first report of a giant virus more than two decades ago (39). Giant viruses are nucleocytoplasmic large double stranded-DNA viruses characterized for their large genome size of at least 200 kb and up to several megabases (40, 41). Giant viruses have been discovered by their co-cultivation with amoebae and only recently have been discovered in diverse environments through metagenomic and single-cell genomic studies (42–44). These studies have enabled us to identify an additional VKOR present in fully sequenced giant viruses. We codon-optimized and cloned five viral *vkor* genes and probed their orientation in the membrane using *Escherichia coli*. We found mutations that adopt the outward configuration and complement disulfide bond formation in the *E. coli* periplasm. We identified a novel truncated γ-carboxylase-like epoxidase fused to a lipid desaturase adjacent in two of the viral VKORs. We performed transcriptomic and proteomic profiling of *Vermamoeba vermiformis* independently infected with *Fadolivirus* and *Yasminevirus* to characterize these vitamin K redox modules during an amoebal infection. We found that both *Fadolivirus* and *Yasminevirus* VKORs and γ-carboxylase-like epoxidase-desaturases are highly expressed during infection throughout the course of infection. We propose that viral VKOR may be involved in regenerating VKH_2_ to drive lipid desaturation and enable the production of the lipid reservoir needed for virion assembly.

## Results

### VKOR homologs in giant DNA viruses have more than one origin

We found viral VKOR when sorting the protein sequences according to the organism’s taxonomy in NCBI. We identified six VKOR sequences in viruses from the Mimiviridae family, 491 sequences in Archaea, 10,213 in Bacteria, 2,013 in Eukaryota and 117 unclassified. These six viral VKOR proteins belong to *Edafosvirus* sp., *Barrevirus* sp. and *Harvfovirus* sp. detected in soil metagenome (42), one VKOR in the viral metagenome (43), and two isolated viruses: *Fadolivirus algeromassiliense* FV1/VV64 from Sidi Bel Abbes (Algeria) sewage water sample (45, 46), and *Yasminevirus saudimassiliense* A1/GU-2018 isolated from Jeddah (Saudi Arabia) sewage water sample (IHU-Mediterranee Infection) (47).

To investigate the evolutionary origin of viral VKORs, we generated a phylogenetic tree using a representative subset of 100 protein sequences from the VKOR domain superfamily (IPR038354). VKOR proteins from *Barrevirus* (Bs)*, Fadolivirus* (Fa)*, Harvfovirus* (Hv) and *Edafosvirus* (Es) clustered together with the VKOR from *Arcella intermedia* rather than with other amoebal VKOR homologs (Fig. 1 cd). In contrast, the *Yasminevirus* (Ys) protein grouped with the archaeal inverted VKOR (VKORi) from *Pyrobaculum aerophilum* (26) (Fig. 1cd). Notably, the commonly used laboratory host *Acanthamoeba* encodes a VKOR homolog that does not cluster with the viral VKORs, whereas *Vermamoeba,* another widely used giant virus model, does not seem to harbor a VKOR. The natural hosts of these giant viruses remain largely unknown, although some are beginning to be elucidated (48). Protein sequence alignment of the six viral VKOR enzymes revealed 22-56% identity (Fig. 2a). The catalytic Cys-XX-Cys (CXXC) motif, required for quinone binding, is conserved across all sequences. Similarly, the CXnC motif, which is essential for interaction with the disulfide carrier, is conserved with a seven-residue spacer in all viruses except *Yasminevirus*, which contains a single-residue spacer (Fig. 2a).

**Figure 2.**
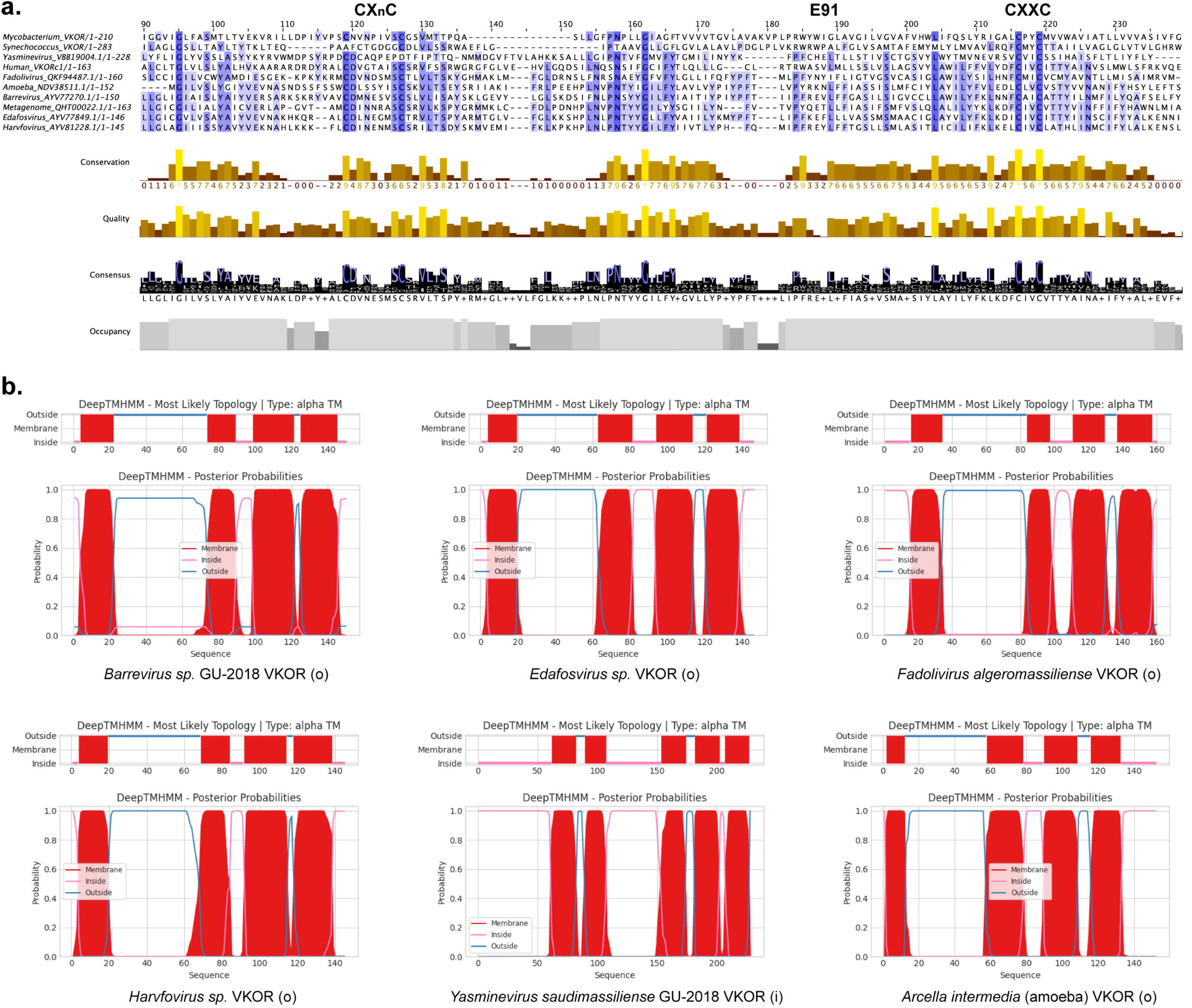
Conservation and topology of viral VKOR proteins. **a.** Multiple sequence alignment of viral proteins shows 22-56% conservation with conserved catalytic residues including the CXXC and CXnC motifs. E91 residue identified in this work is also indicated. Alignment was done using Clustal Omega (https://www.ebi.ac.uk/jdispatcher/msa/clustalo) (92) and visualization using Jalview 2.11.4.1 (93). **b.** Topology prediction plots suggest an outward orientation for all except *Yasminevirus saudimassiliense*. VKOR. Predictions were done using Deep TMHMM (https://dtu.biolib.com/DeepTMHMM) (51). o: cysteines facing outward, i: cysteines facing inward.

These findings suggest that viral VKOR proteins may have originated from different ancestors or perhaps transferred from an ancient unknown host particularly acquired by ancient lateral gene transfer. Notably, Megaviricetes are known to show horizontal gene transfer with amoeba and multicellular animals (43). However, expanding viral and natural hosts genome sequencing efforts could help clarify viral VKOR evolutionary history.

### Membrane topology analysis in E. coli suggests viral VKOR outward orientation

Members of the Crenarchaea, such as *Aeropyrum pernix*, have two VKOR paralogs with differing orientations in the membrane, one that orients the four catalytic cysteines outward into the extracytosolic space (*Ap*VKORo), while a second orients the catalytic cysteines inward to the cytoplasm (*Ap*VKORi) (26). The rest of the VKOR orthologs (bacterial, plantae and mammalian) adopt the outward configuration (20, 34). We determined the predicted membrane topology of viral VKOR proteins using TOPCONS, which uses a consensus of predictions from several algorithms (49, 50) (Supplementary Fig. 1) and Deep TMHMM, which uses deep learning model and overall outperforms in accuracy other algorithms (51) (Fig. 2b). We found mixed topology predictions for VKOR proteins using TOPCONS (Supplementary Fig. 1), however, they are predominantly predicted to face out using Deep TMHMM except for *Ys*VKOR (Fig. 2b). Indeed, *Ys*VKOR clustered with *P. aerophilum* VKORi (Fig. 1d).

We used *E. coli* to determine the functionality and orientation of these five viral VKOR proteins (Fig. 3a). The study of diverse VKOR catalysts in *E. coli* is possible given that VKOR homologs from actinobacteria, crenarchaea and three eukaryotes (plant, rat and human) can complement periplasmic disulfide bond formation of *E. coli* lacking *dsbB* despite the lack of homology between VKOR and the native DsbB enzyme (18, 26, 28, 29, 52, 53). Thus, disulfide exchange reactions are widely conserved and can be studied using *E. coli* motility assays. That is, the VKOR enzyme facing outward would oxidize *E. coli* DsbA which oxidizes the flagellar P-ring protein, FlgI, restoring motility of the *E. coli* Δ*dsbB* mutant. Alternatively, if the VKOR is oriented towards the cytoplasm, we used an *E. coli* strain containing a plasmid encoding a signal sequence-less alkaline phosphatase (Δss*phoA*), which localizes to the cytoplasm (54). Since disulfide bond formation ordinarily does not occur in the *E. coli* cytoplasm, the alkaline phosphatase does not fold properly and is enzymatically inactive. However, this enzyme can be activated by reconstituting the disulfide bond formation pathway in the cytoplasm (26). That is, by expressing a signal sequence-less *E. coli* DsbA and a VKOR facing inward, such as archaeal *Ap*VKORi (26). Hence, under these conditions cytoplasmic alkaline phosphatase acquires its disulfide bonds and becomes active. This activity can be detected by the ability of cytoplasmic alkaline phosphatase to replace other cytoplasmic phosphatase, serine-1-phosphatase (SerB) (54). Thus, while a strain lacking *serB* is auxotrophic for serine, if alkaline phosphatase is active in the cytoplasm, it can replace the missing biosynthetic phosphatase and restore growth in the absence of serine.

**Figure 3.**
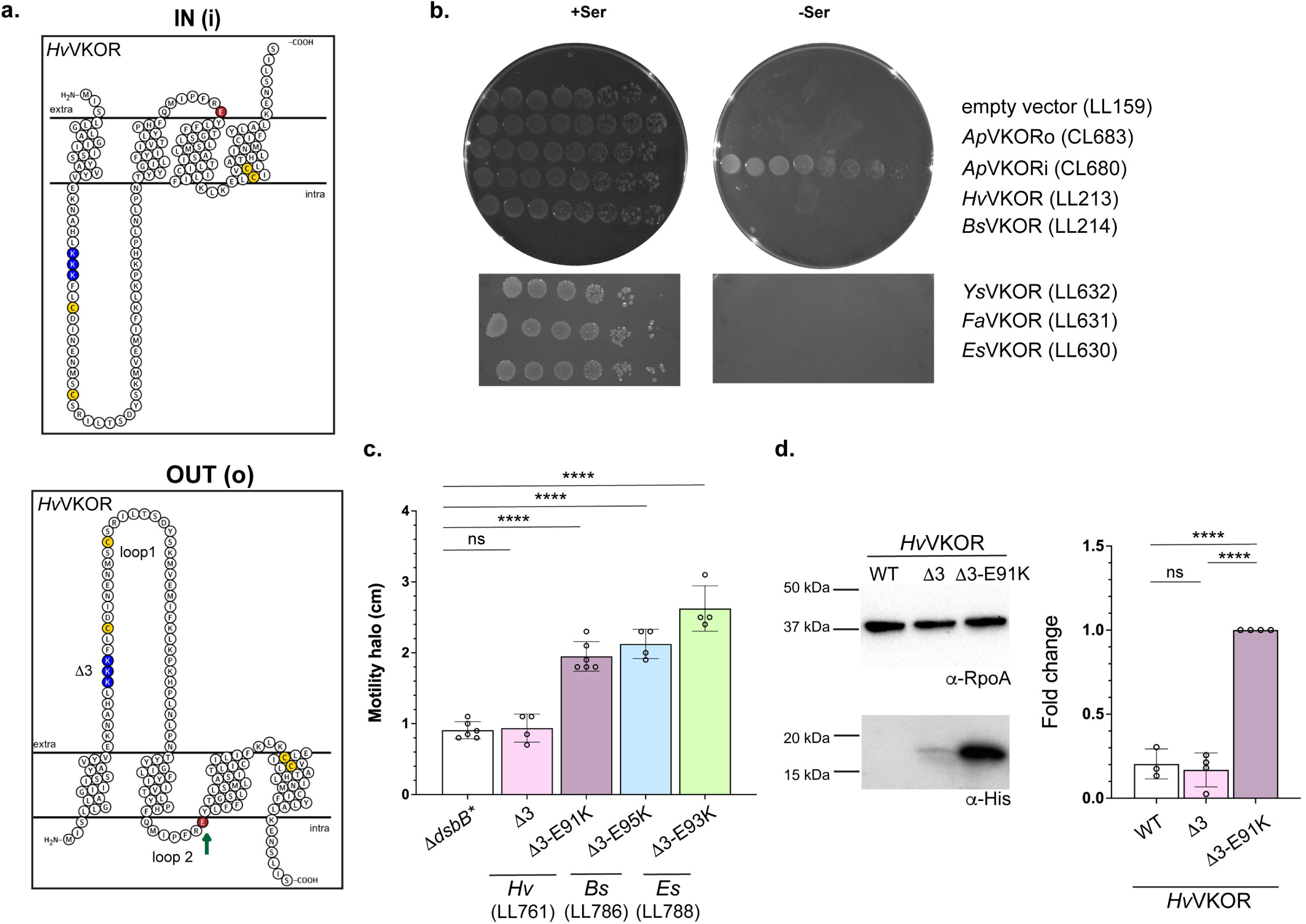
Viral VKORs adopt an outward orientation in *E. coli*. **a**. *Harvfovirus* sp. VKOR (*Hv*VKOR, UniProt A0A3G5A1Y0) is shown as an example of inward (top) and outward (bottom) topology. Topology visualizations were done using Protter (83). Catalytic cysteines are highlighted in yellow, while positive and negative residues are highlighted in blue and red, respectively. **b**. *E. coli* serine auxotrophy assay to test cytoplasmic orientation of catalytic cysteines. Strains were grown on M63 minimal media with or without serine. Plates were incubated for 2 days at 37°C. **c**. *E. coli* motility assay to test periplasmic orientation of catalytic cysteines. Random mutagenesis produced a mutant (E91K) that conferred motility that adopts an outward orientation. Balancing the positive residues in the first two loops in *Barrevirus* sp. VKOR and *Edafosvirus* sp. VKOR also adopt an outward topology in *E. coli*. Motility halos were measured after 3 days of incubation at 30°C, concentrations of IPTG used are 25 µM for *Hv*VKOR, 10 µM for *Bs*VKOR and 50 µM for *Es*VKOR. **d**. Removal of positive residues in the extracytoplasmic loop together with addition of a positive charge in the cytoplasmic loop stabilize *Hv*VKOR. Protein abundance was quantified using anti-His antibody and normalized using anti-RpoA antibody as a loading control. Data represents average±SD of at least three biological replicas. Statistical tests were done using one-way ANOVA multiple comparisons. p-values are depicted in GP style: p-value ≤0.0001 (****), 0.0002 (***), 0.021 (**), 0.0332 (*). Non-significant p-values (>0.1234) were indicated as ns.

We codon optimized and cloned five viral *vkor* genes in a plasmid under an IPTG-inducible promoter, which also harbors the signal sequence-less *E. coli dsbA*. We then independently transformed the plasmids into two *E. coli* strains: Δ*serB*Δss*phoA* (DHB7787 strain) for testing the inward orientation and Δ*dsbB* mutant (HK325 strain) for outward orientation. For the inward orientation of the five viral VKORs, we observed that while our positive control expressing archaeal *Ap*VKORi grows in the absence of serine, none of the strains harboring viral VKORs grew in the absence of serine (Fig. 3b). Thus, suggesting that viral VKORs may not be able to adopt the inward orientation in the *E. coli* membrane. When we tested the outward orientation by transforming the plasmids into a Δ*dsbB* mutant, we similarly observed that none of them complemented the motility defect of *dsbB* mutant (Supplementary Fig. 2).

We reasoned that viral VKORs are likely inserted into the ER membrane, which could explain the lack of complementation in *E. coli*. Therefore, we instead used a strain that allows proper insertion of foreign membrane proteins into the *E. coli* membrane (25). Mammalian VKORc1 homologs can complement a *dsbB* mutant through two changes: (1) the deletion of positive residues in the first extracytoplasmic loop of the protein and (2) two additional chromosomal mutations that modify the insertase YidC (T362I) and inactivate the protease HslV (C160Y) (25, 53). Presumably, this combination of mutations altogether prevents degradation of the VKORc1 and allows its proper insertion. To test the viral genes, we transformed plasmids harboring viral *vkor* genes into a Δ*dsbB* mutant carrying the *yidC*T362I *hslV*C160Y alleles (Δ*dsbB\**). We also deleted three positive residues from extracytoplasmic loop 1 of *Barrevirus* (ΔR27K28K30, from here on *Bs*VKORΔ3) and *Harvfovirus* (ΔK26K27K29, from here on *Hv*VKORΔ3) proteins and tested their activity in the enhanced ER membrane insertion background (Δ*dsbB\**).

Although none of these constructs complemented the motility defect of *dsbB* mutant (Supplementary Figs. 2 and 3), we observed that some *E. coli* strains developed motility halos after three days of incubation with wildtype *Barrevirus*, *Harvfovirus* and *Yasminevirus* VKOR proteins (Supplementary Fig. 4). However, plasmids isolated from these suppressors did not restore motility when re-transformed into the Δ*dsbB* mutants (Supplementary Fig. 4), indicating that the suppressor mutations occurred in the *E. coli* chromosome. Genome sequencing revealed the chromosomal mutations, three out of four mapped to outer membrane protein A (one isolate had OmpAL222fs combined with the cell wall hydrolase NlpCH115R, and two isolates carried OmpAR328L and OmpAG234fs, respectively). The OmpA mutations disrupt the protein or the OmpA-like domain, while the NlpC mutation disrupts one of the catalytic histidine residues known for its role as proton acceptor in the hydrolase domain. *E. coli* strains lacking OmpA have been previously reported to display increased motility, presumably because OmpA is a major substrate of the Dsb machinery and therefore a drain of disulfide bonds (55). The fourth mutant carried a change in the two-component membrane associated histidine kinase, PhoQ (PhoQL387Q), disrupting leucine 387 which is part of the ATP binding site thus disrupting the phosphorelay system. Increased expression of PhoP-regulated genes via PhoQ has been previously reported in *dsbA* or *dsbB* mutants (56). Altogether, these mutations presumably restore motility by decreasing disulfide bond formation in abundant *E. coli* proteins or regulate the expression of genes that alter surface components.

Since the mutations do not relate to the insertion of the viral VKOR proteins into *E. coli* membrane, we next performed random mutagenesis of three viral *vkor* genes (*Ysvkor*, *Hvvkor* and *Hvvkor*Δ3) and selected for mutants that could restore either *E. coli* motility (outward) or serine auxotrophy (inward) to probe both topology and functionality. We generated plasmid libraries containing approximately ∼30,000 independent mutants in each library and transformed them into two strains, Δ*serB*Δss*phoA* and Δ*dsbB\**. We then plated the libraries either on minimal media lacking serine or into motility plates, respectively. We only obtained one motility halo for the Δ*dsbB*Hv*VKORΔ3 library and no colonies on minimal media lacking serine for any of the libraries. Suppressor plasmid isolation and re-transformation confirmed that it conferred motility to the Δ*dsbB\** mutant (Fig. 3c). Plasmid sequencing identified a single mutation in *Hv*VKOR, E91K, which converts a negative residue into a positive one in the second loop predicted to be cytoplasmic (Fig. 3a). This suggests that viral VKORs, similar to human and rat VKORs, violate the positive-inside rule and that their insertion into the *E. coli* membrane requires a balance of positive charges between periplasmic and cytoplasmic loops (52, 53).

To determine if the combination of removing three positive residues in loop 1 and the addition of a positive residue in loop 2 stabilizes the protein, we assessed protein abundance in cell lysates using an anti-His antibody to detect *Hv*VKOR. Both wildtype and *Hv*VKORΔ3 accumulate at low levels or below the limit of detection in Δ*dsbB\**, whereas the combined Δ3-E91K mutant has six-fold higher protein levels (Fig. 3d). These results indicate that the positive residue in loop 2 stabilizes the protein when combined with removal of three positive residues in loop 1.

To determine whether similar changes could help other viral VKORs complement *E. coli* Δ*dsbB*, we introduced a glutamate-to-lysine substitution in *Bs*VKOR (E95K) and *Es*VKOR (E93K), together with deletion of three positive residues in loop 1 (R27K28K30 and K24K26R28, respectively). We then tested whether these changes conferred motility to Δ*dsbB\** strain. Both mutated VKORs were able to adopt the outward topology and complement motility in *E. coli* with these changes (Fig. 3c). Thus, these three viral VKORs may adopt the outward insertion in the ER membrane.

### Viral VKOR is adjacent to a truncated vitamin K γ-carboxylase fused to a desaturase

We analyzed the genetic context of the five VKORs in their assembled viral genomes and found an adjacent vitamin K-dependent γ-carboxylase-like epoxidase domain (from here on VKE for vitamin K epoxidase) in all cases except *Edafosvirus* (Fig. 5a). Human VKGC (*Hs*VKGC) is an integral membrane protein of the ER that catalyzes the conversion of glutamic acid to γ-carboxyglutamic acid (Gla) (57, 58). This post-translational modification activates the blood-clotting cascade in vitamin K-dependent coagulation factors (58). In *Harvfovirus* and *Yasminevirus*, the carboxylase is fused to a sterol desaturase gene, forming a single protein (from here on VKED for vitamin K epoxidase desaturase), whereas in *Barrevirus* and *Fadolivirus*, the sterol desaturase gene is positioned in the opposite orientation (Fig. 4a). An alignment between the viral γ-carboxylase-like epoxidases and the *Hs*VKGC revealed that although the viral enzymes retain the catalytic lysine (K218) required for deprotonation and epoxidation of VKH_2_ (Fig. 4c), they lack residues 374-418 which are highly conserved across vertebrate and invertebrate species and essential for γ-carboxylation (Fig. 4c) (59, 60). Inspection of the surrounding viral genome region revealed no small ORFs with carboxylase signatures, consistent with the absence of these residues. In line with this observation, we were unable to find a substrate for the putative γ-carboxylases by searching for the Gla domains in the *Barrevirus* and *Harvfovirus* proteomes (PDOC00011 domain search in PROSITE Expasy) (61).

**Figure 4.**
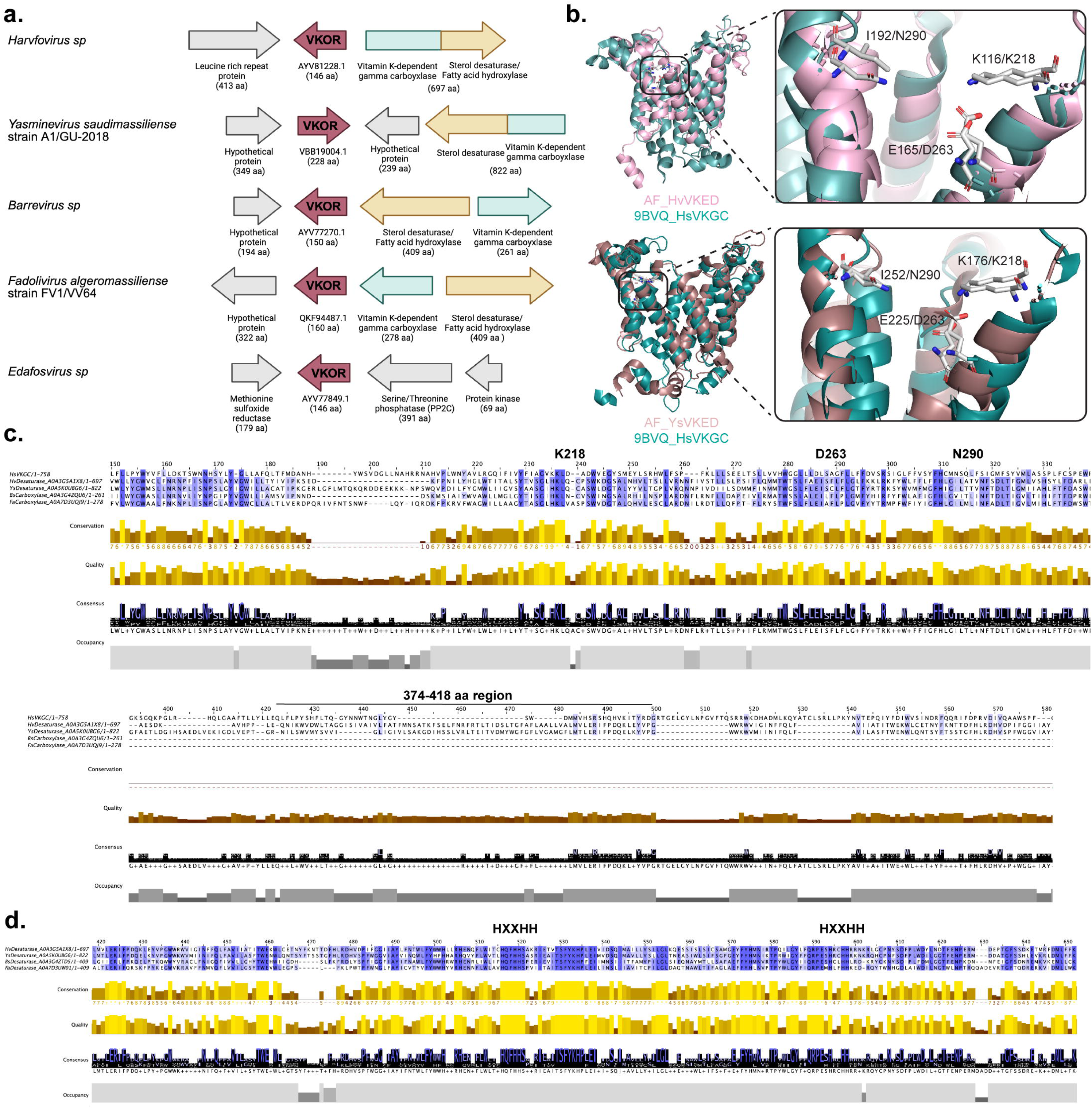
Vitamin K γ-carboxylase-like epoxidase adjacent to viral VKOR proteins. **a**. Genetic context of *vkor* loci (red) in five giant viruses. All virus except *Edafosvirus* (*Es*) harbor a vitamin K dependent γ-carboxylase (green) and a sterol desaturase (yellow) adjacent to VKOR. These two proteins are fused forming a single polypeptide in *Harvfovirus* (*Hv*) and *Yasminevirus* (*Ys*). **b**. Viruses harbor a truncated vitamin K carboxylase. Top: Superimposition of residues 1-230 of *Harvfovirus* sp. vitamin K epoxidase-desaturase*, Hv*VKED (A0A3G5A1X8. Structure obtained with AlphaFold3 (94) pLDDT 81.8) and residues 1-321 of human VKGC (9BVQ) (63) using PyMOL. Root mean square deviation value (RMSD) of superimposition is 2.63. Bottom: Superimposition of residues 1-300 of *Yasminevirus* vitamin K epoxidase-desaturase*, Ys*VKED (A0A5K0UBG6. Structure obtained from AlphaFoldDB [https://alphafold.ebi.ac.uk/] pLDDT 84.01) and residues 1-321 of human VKGC (9BVQ) (63) using PyMOL. Root mean square deviation value (RMSD) of superimposition is 2.47. Catalytic residues are indicated for *Hv*VKED, *Ys*VKED and *Hs*VKGC, respectively. **c**. Protein alignment of vitamin K γ-carboxylases showing conservation of catalytic residue (K218), but the lack of conserved residues 384-415 found in human γ-carboxylase and vertebrate/invertebrate γ-carboxylases. Residues D263 and N290 at the catalytic center are also indicated. **d**. Protein alignment of lipid desaturases showing conservation of the two HXXHH motifs that coordinate two iron molecules and form the catalytic center.

**Figure 5.**
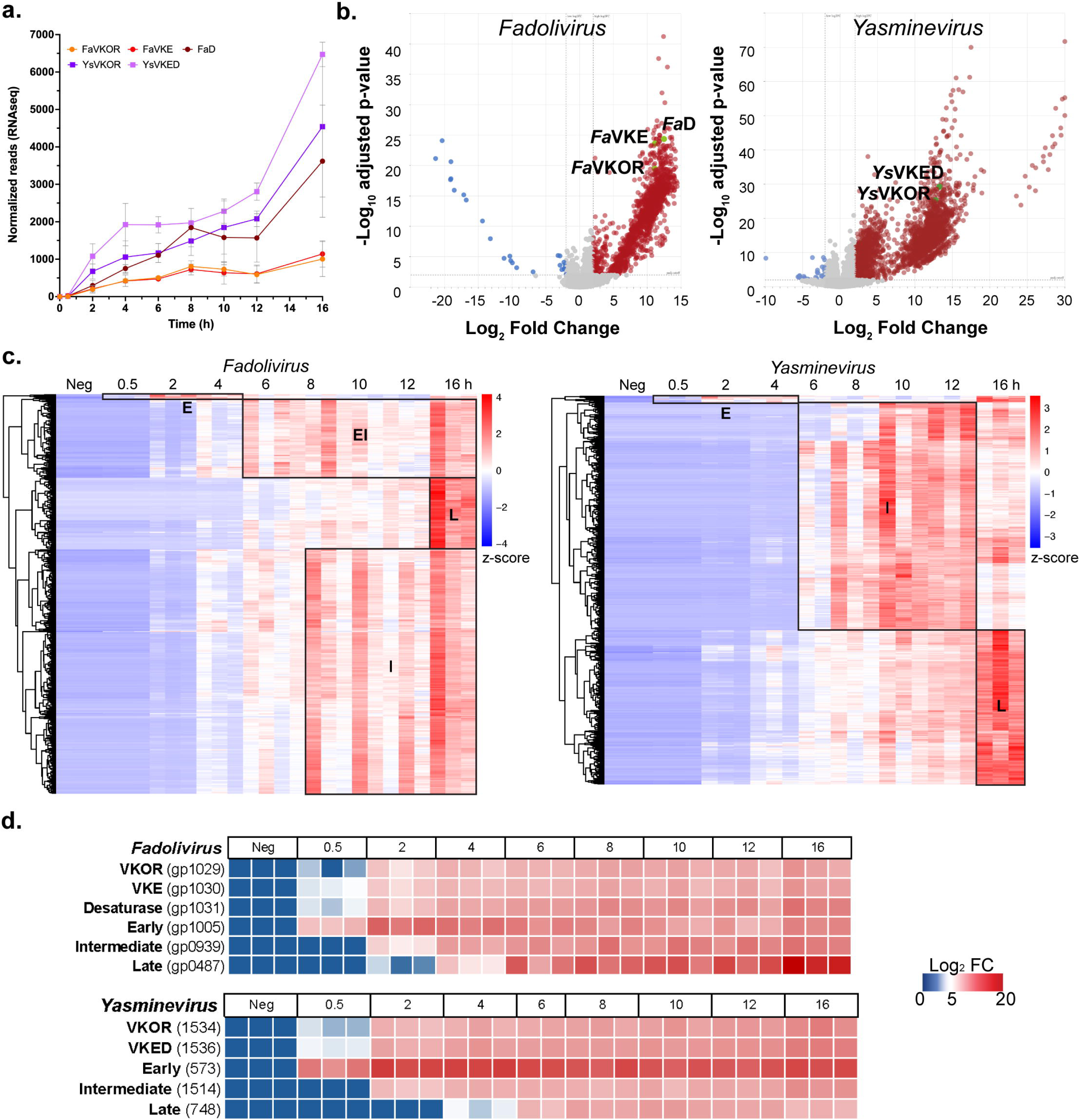
Vitamin K redox modules are abundantly transcribed as intermediate genes during *Fadolivirus* and *Yasminevirus* infections of *V. vermiformis*. **a**. RNA normalized reads detected in samples infected with *Fadolivirus* (*Fa*) or *Yasminevirus* (*Ys*) compared to the mock infected control (T0). **b**. Volcano plots of differentially expressed viral genes compared to the mock infected control. Down (blue) and upregulated (red) genes are indicated. Vitamin K redox modules are highlighted in green, including: *Fa*VKE: vitamin K epoxidase; *Fa*D: Desaturase; *Fa*VKOR: vitamin K epoxide reductase; *Ys*VKED: vitamin K epoxidase desaturase; and *Ys*VKOR: vitamin K epoxide reductase. **c**. Gene expression clustering of *Fadolivirus* or *Yasminevirus* genes when infecting *V. vermiformis* through a 16 h course of infection. Neg, mock infected. Data represents z-score of each of the three independent replicas. **d**. Comparison of gene expression patterns of early, intermediate and late genes with Vitamin K redox modules. Neg, mock infected. Data represents log_2_ fold change of each of the three independent replicas.

The recently solved structure of *Hs*VKGC has elucidated how epoxidation is coupled to γ-carboxylation (62, 63). To compare the viral γ-carboxylases with *Hs*VKGC, we used AlphaFold3 to predict the structure of *Harvfovirus* VKED (*Hv*VKED) and superimposed it on *Hs*VKGC structure. We were able to align the first 230 residues of *Hv*VKED or the first 300 residues of *Ys*VKED to the first 300 residues of *Hs*VKGC, which encompass the epoxidase domain yielding RMSDs of 2.63 and 2.47, respectively (Fig. 4b). In *Hs*VKGC, the hydroxyl groups of VKH_2_ form hydrogen bonds with N290 (*Hv*VKED-I192 and *Ys*VKED-I252) and the D263-K218 pair (Fig. 4b) (63). Epoxidation is initiated by K218 (*Hv*VKED-K116 and *Ys*VKED-K176), which deprotonates the 4-hydroxyl group of VKH_2_, a reaction facilitated by D263 (*Hv*VKED-E165 and *Ys*VKED-E225) (63). Notably, D263 aligns to E165/E225 in the viral proteins (Fig. 4b), and mutational studies have shown that *Hs*VKGC variants at D263 lose activity except for D263E, which retains 25% of activity (63). Together, these results suggest that the viral γ-carboxylases are truncated, retaining only the VKE domain.

Given these findings, we hypothesize that the annotated γ-carboxylase-like epoxidase performs an alternative function in viruses. In *Harvfovirus* and *Yasminevirus*, the fusion of the epoxidase domain with a sterol desaturase, suggests a potential role in lipid biosynthesis. Members of the desaturase family are integral ER membrane proteins involved in cholesterol biosynthesis and contain two cytoplasmic HXXHH motifs that coordinate two iron molecules at the catalytic center. Alignment of the viral desaturases adjacent to VKOR from *Harvfovirus, Barrevirus*, *Fadolivirus* and *Yasminevirus* confirmed that all four contain the two cytoplasmic HXXHH motifs (Fig. 4d), thus altogether suggesting that they could catalyze fatty acid desaturation coupled to vitamin K epoxidation (see Discussion, Fig. 6).

**Figure 6.**
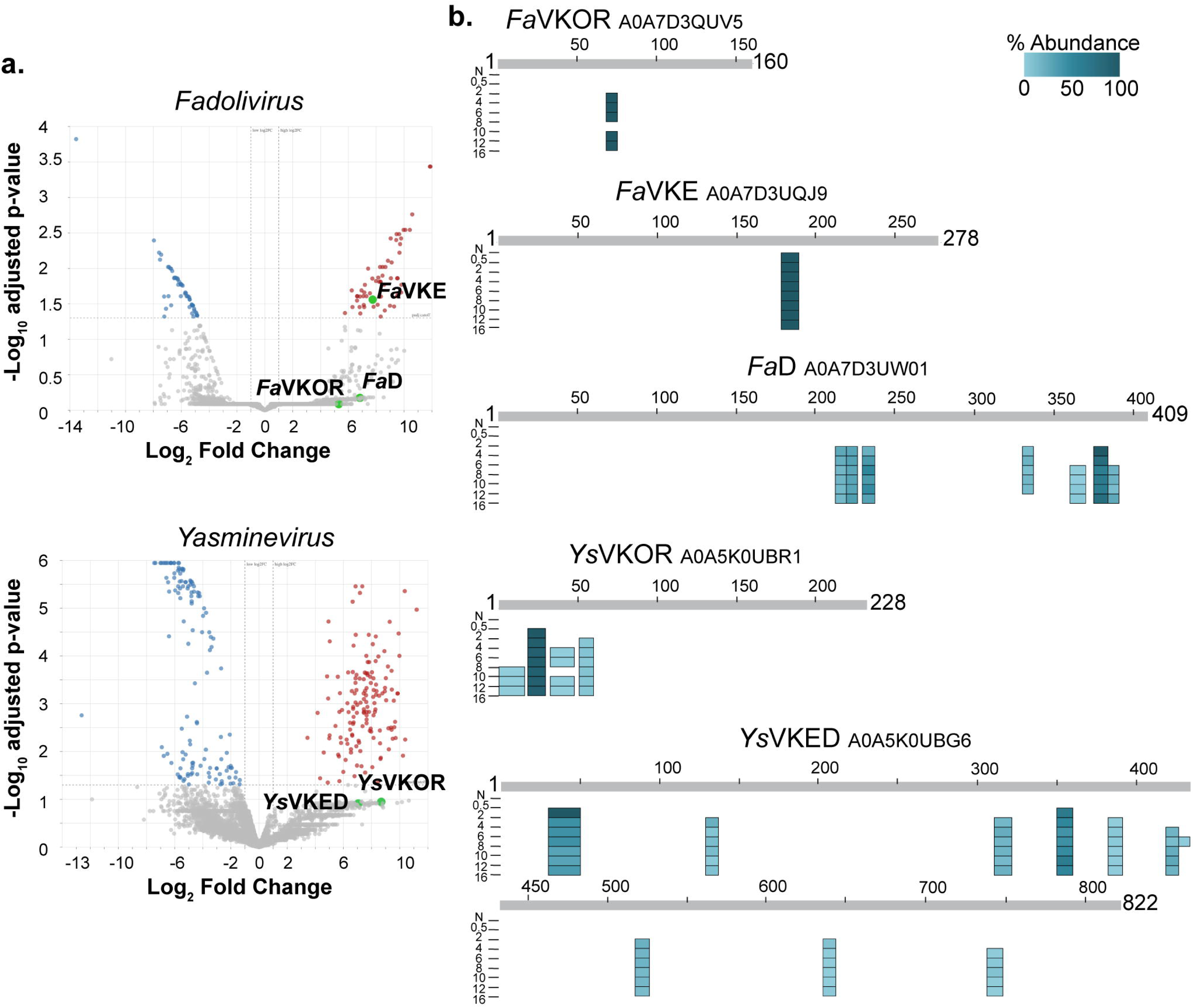
Vitamin K redox proteins are detected after 2-4 h of *Fadolivirus* (*Fa*) and *Yasminevirus* (*Ys*) infections of *V. vermiformis*. **a**. Volcano plots of viral protein abundance compared to the mock infected control through a 16 h course of infection. Low (blue) and high (red) abundance proteins are indicated. Vitamin K redox modules are highlighted in green, including: *Fa*VKE: vitamin K epoxidase; *Fa*D: Desaturase; *Fa*VKOR: vitamin K epoxide reductase; *Ys*VKED: vitamin K epoxidase desaturase; and *Ys*VKOR: vitamin K epoxide reductase. **b**. Peptide coverage of *Yasminevirus* and *Fadolivirus* Vitamin K redox proteins through a 16 h course of infection. Peptide abundance was normalized to the total peptides abundance and indicated as percentage (%). N, mock infected control.

### Intermediate-stage transcription and translation of viral vitamin K redox modules during Vermamoeba vermiformis infection

To start characterizing viral VKOR, vitamin K-dependent γ-carboxylase-like epoxidase (VKE) and desaturase (VKED or D) we used two previously developed giant virus infection models using *Fadolivirus algeromassiliense* FV1/VV64 and *Yasminevirus saudimassiliense* A1/GU-2018 (45, 47). We infected *V. vermiformis* strain CDC19 with *Fadolivirus* or *Yasminevirus* followed by RNAseq and proteomic analyses.

The transcriptional profiling was performed at 0.5, 2, 4, 6, 8 10, 12 and 16 h post-infection. Poly-adenylated RNA was analyzed by RNA-Seq using a mock-infected culture as negative control. Three biological replicates for each condition were sequenced using the Illumina NovaSeq platform. Quality control analyses and read alignment data are shown in Supplementary Fig. 5 and Supplementary Table 4a. The mapped reads to *Fadolivirus* and *Yasminevirus* genomes displayed a higher percentage of reads mapped at or after H6, which is typical for a giant virus course of infection (64). The DESeq2 pipeline was used to identify differentially expressed genes in the infected amoebae at the different times post-infection compared to mock infected controls. The gene lists where –log_10_ adjusted p-value ≥ 2 and log_2_ fold change ≥ 2 and the up and down-regulated genes are provided in Supplementary Table 4bc. We were able to detect transcripts of all five genes: *Ys*VKOR (Yasminevirus_1534), *Ys*VKED (Yasminevirus_1536), *Fa*VKOR (QKU48_gp1029), *Fa*VKE (QKU48_gp1030) and *Fa*D (QKU48_gp1031), and their expression increased over time (Fig. 5a). We found that *Ys*VKOR and *Ys*VKED resulted in a 12.7 and 13.2 log_2_ fold change compared to the mock infected control, while *Fa*VKOR, *Fa*VKE and *Fa*D in a 11.1, 11.1, and 12.5 log*2* fold changes, respectively (Fig. 5b, adjusted p-values <10^-20^). Thus, these vitamin K modules were highly expressed throughout the viral infections.

We performed hierarchical gene clustering into groups sharing similar profiles based on their expression levels at eight time points (Fig. 5c and Supplementary Table 4de). For *Fadolivirus* the clusters reflect three temporal expression classes that coincide with the traditional classification of viral genes into ‘‘early’’ (E), expressed immediately after 30 min and until 4 h after infection; ‘‘Intermediate’’ (I) genes which were clustered into two subgroups: early intermediate (EI) transcribed after 4 h and until 16 h and intermediate (I) transcribed between 8 and 16 h. ‘‘Late’’ (L) genes which were transcribed mostly after 12-16 h (Fig. 5c). Similarly, for *Yasminevirus* we found early genes from 30 min to 6 h; intermediate from 8 to 12 h; and late majorly expressed until 16 h (Fig. 5c). When we compared the vitamin K module expression patterns to those of early, intermediate and late genes, we observe that *Fa*VKOR, *Fa*VKE, *Fa*D, *Ys*VKOR and *Ys*VKED are transcribed similar to intermediate genes (Fig. 5d). Additionally, the distribution of functional categories of the expressed genes shows that genes associated to metabolic functions, such as these vitamin K redox genes, increase after 2 h post-infection and are maintained until 16 h in both *Fadolivirus* and *Yasminevirus* infections (Supplementary Fig. 6). Collectively, the transcriptional profiling suggests that vitamin K redox genes contribute at an intermediate phase of infection to alter the host’s metabolism.

Lastly, we analyzed the proteome of *V. vermiformis* samples infected with either *Fadolivirus* (Supplementary Table 5 and 7) or *Yasminevirus* (Supplementary Table 6 and 8) during the same course of infection. We identified 1,028 proteins of *Fadolivirus* while 997 of *Yasminevirus* together with 6,718 and 6,587 proteins from *V. vermiformis*, respectively (Supplementary Table 9). We were able to identify peptides of *Fa*VKOR, *Fa*VKE, *Fa*D in *Fadolivirus*-infected samples, while *Ys*VKOR and *Ys*VKED proteins in *Yasminevirus-*infected samples after 2 h post-infection of amoebae but none in the mock infected control. Their protein abundances ranged between 5-8 log_2_ fold change but did not meet the log_10_ adjusted p-value cutoff of -1.3, except for *Fa*VKE (Fig. 6a). The low peptide abundance and coverage (Fig. 6b) may reflect technical limitations such as poor solubility of the membrane proteins, poor ionization efficiency, or limited tryptic peptide detectability likely reducing the statistical power to confidently detect significant changes. Notably, however, the peptide coverage of the membrane protein *Ys*VKED includes peptides from both the epoxidase and desaturase domains (Fig. 6b) and this is consistent with the RNA reads of the gene transcribed as a single open reading frame (Supplementary Fig. 7). Altogether our transcriptomic and proteomic data suggest that the viral VKOR and epoxidase-desaturases contribute to modulate an effective giant virus infection.

## Discussion

Most VKOR proteins orient their catalytic centers toward the extracytoplasmic space, with the exception of some crenarchaeal VKORs (25, 26). The study of diverse disulfide bond catalysts in *E. coli* has been possible despite the lack of homology between VKOR and DsbB proteins. VKOR homologs from actinobacteria (*Mycobacterium tuberculosis*) (18), archaea (*A. pernix*) (26), firmicutes (*Clostridium sp.*) (65), and three eukaryotes (plant, human and rat) (28, 52, 53) can complement disulfide bond formation in *E. coli*. Here, we have discovered viral VKOR homologs and expressed them functional in *E. coli* by removing three positively charged residues in the first extracytoplasmic loop and substituting a negatively charged residue with a positive one in the first cytoplasmic loop. Mammalian and viral VKORs are ER membrane proteins, thus, require modifications that balance the positive charges in the first two loops to stabilize viral VKOR and appropriately insert into the *E. coli* membrane. Once inserted into the membrane, the viral VKORs can participate in disulfide exchange reactions with *E. coli* DsbA and transfer electrons to ubiquinone-8, the predominant quinone under aerobic growth (15). We did not explore additional modifications of *Fadolivirus* and *Yasminevirus* VKORs because they lack a negative residue aligned to Glu91 (Asn and His, respectively). Moreover, *Yasminevirus* VKOR contains a CXC rather than a CXnC motif, which may limit its interaction with *E. coli* DsbA. These proteins may therefore require further modifications in the first and second loops to insert in the *E. coli* membrane and interact with *E. coli* DsbA effectively either in the cytoplasm (*Yasminevirus*) or periplasm (*Fadolivirus*). Our studies suggest that all except *Yasminevirus* viral VKOR homologs may adopt outward insertion in the host ER.

Our transcriptomic and proteomic profiling identified transcripts and peptides of VKOR and the vitamin K-dependent γ-carboxylase-like epoxidases/desaturases in both *Yasminevirus* and *Fadolivirus* infecting *V. vermiformis* thus suggesting a major role during the viral infection. Phospholipids are essential for maintaining the structure, function, and stability of cell membranes. Their distribution within the bilayer determines curvature, fluidity, permeability, and the interactions with membrane-associated protein complexes (66). Proper membrane packing and fluidity are critical for replication of giant viruses. In contrast to most known DNA viruses, most of large DNA viruses do not deliver their genome to the nucleus but instead replicate and assemble in cytoplasmic viral factories (Fig. 7a) (67). Therefore, a critical process during a giant virus infection is the ER-recruitment to the periphery of the viral factory which is followed by *de novo* synthesis of lipid precursors that generate the internal single layer lipid membrane that surrounds the virus core (Fig. 7a) (67–69). This lipid membrane would then fuse with the vacuole membrane to release the double-stranded DNA (Fig. 7a) (68).

**Figure 7.**
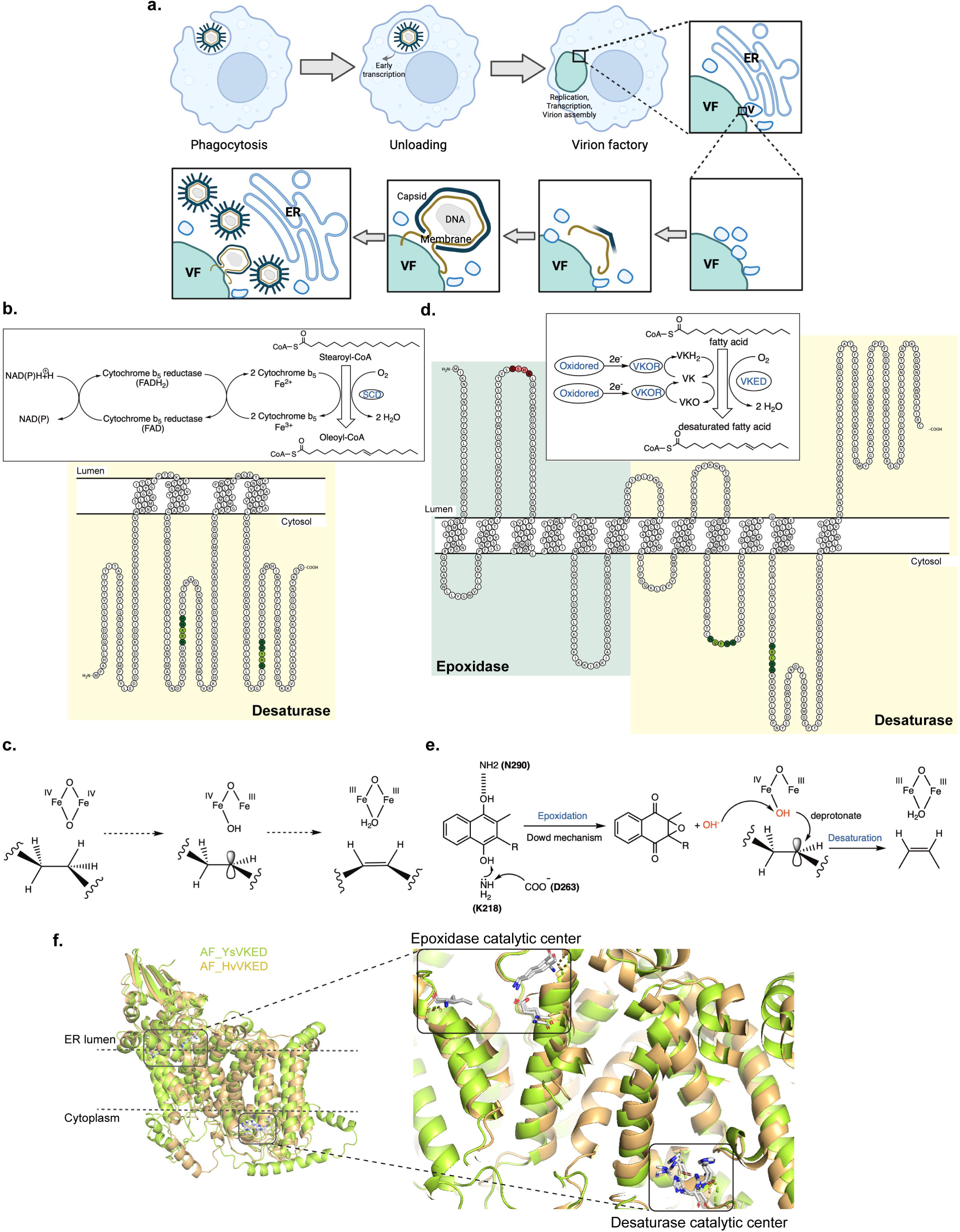
Epoxidation coupled to lipid desaturation in giant viruses. **a**. Giant virus infection model. Virus is phagocytosed by amoeba, unloads its genome and early transcription starts. A cytoplasmic virion factory is assembled where genome replication; intermediate and late transcription; as well as membrane and capsid assembly take place. Host ER cisternae are recruited to the viral factory (VF) and produce small vesicles (V). Multivesicular bodies form membrane sheets that generate the viral membrane monolayer that together with the capsid produce the icosahedral morphology. Host ER cisternae act as a lipid reservoir that enables continuous viral assembly. Adapted from (69, 95) **b**. Desaturation of stearoyl-CoA by human SCD. Adapted from (96). The membrane topology is shown for human SCD1 (Uniprot O00767). **c**. Generic mechanism of desaturation. The structures of the di-iron oxidant and the reactive intermediates are speculative. Adapted from (71). **d**. Proposed mechanism of the viral vitamin K epoxidase desaturase (VKED). This enzyme could couple epoxidation to drive lipid desaturation by using VKO to generate the carbon-centered radical/FeOH pair to then produce the double bond and iron-bound water. The membrane topology is shown for *Hv*VKED. **e**. Proposed coupling mechanism of epoxidation with lipid desaturation based on the coupling mechanism of epoxidation and γ-carboxylation by *Hs*VKGC. Adapted from (63). **f**. Superimposition of AlphaFold (94) predicted structures of *Hv*VKED (A0A3G5A1X8) and *Ys*VKED (A0A5K0UBG6). Structural alignment was done using PyMOL with RMSD value of 1.85. Catalytic centers are indicated in squares. Topology visualizations were done using Protter (83).

A common strategy to remodel membrane architecture is to alter the saturation and the length of fatty acid hydrophobic tails (70). Unsaturated fatty acids are synthesized in the cytoplasm, where, following an initial desaturation step, they undergo elongation and further desaturation before being incorporated into triglycerides, cholesterol esters, or phospholipids. This initial desaturation is the rate-limiting step in unsaturated fatty acid biosynthesis and is catalyzed by stearoyl-CoA desaturase (SCD, Fig. 7b) (66). SCD introduces a *cis*-double bond at the Δ9 position of stearoyl-CoA and palmitoyl-CoA (66). Located in the ER, SCD uses NADH as a reductant and functions together with the flavoprotein cytochrome b5 reductase, cytochrome b5, and molecular oxygen (Fig. 7b). Human SCD1 contains four transmembrane segments and eight cytoplasmic histidine residues that coordinate iron within the enzyme’s catalytic center also known as di-iron binding motif (Fig. 7b). The reaction is oxygen dependent, and the di-iron hydroxyl abstracts a hydrogen or proton from the fatty acid generating the double bond in the fatty acid while reducing molecular oxygen into water (71). Importantly, desaturation is initiated by an energetically expensive C–H activation step to produce a hypothetical carbon-centered radical/FeOH pair which generates the double bond and iron-bound water (Fig. 7c) (71). The genetic context of the viral VKOR proteins identified in this work suggests that VKOR may act in concert with a vitamin K epoxidase and a lipid desaturase, potentially enabling these viruses to perform lipid desaturation. The four viral desaturases identified near VKOR harbor the eight-histidine motif facing the cytosol as in human SCD1, while the catalytic lysine predicted to participate in epoxidation localizes to the ER lumen (Fig. 7df). We hypothesize that a host luminal oxidoreductase is oxidized by the four viral VKORs, and a cytoplasmic oxidoreductase in the case of *Yasminevirus* VKOR. Then, VKOR uses the two electrons to reduce VKO to VK and then VK to VKH_2_ (Fig. 7d). Subsequently, VKH_2_ provides the electrons to the viral VKED to desaturate a fatty acid. In *Harvfovirus* and *Yasminevirus* the domains would act in concert given the fusion of the two domains (Fig. 7f) while in *Barrevirus* and *Fadolivirus* the two enzymes would interact to catalyze the desaturation step. The reaction could then follow the binding of VKH_2_ to the viral epoxidase domain at the membrane, then the generation of the VKO and a free hydroxide ion, which could then diffuse and promote at the cytoplasmic desaturase domain the carbon radical/FeOH center in the fatty acid (Fig. 7ef). The physical separation of the two catalytic centers is also observed in VKGC, which is resolved by generating a diffusive superbase (OH^-^) (63). Although, both centers face the ER lumen in VKGC, proteins that catalyze electron transfer between spatially separated sites have been previously described (72).

Auxiliary metabolic genes represent a powerful strategy of host metabolism manipulation by expanding the metabolic capabilities of infected cells, particularly in fluctuating and often harsh environments such as those encountered by amoebae. The presence of an alternative lipid modification module that uses a distinct electron source may provide a selective advantage, enabling viral lipid synthesis independently of host NADH source and instead drawing on a vitamin K-dependent electron pool. This vitamin K-based system represents an additional−or potentially complementary−alternative layer of lipid metabolism to that recently described in giant viruses, which includes novel soluble cytochrome b_5_ proteins (73). Together, these findings underscore the importance of lipid biosynthesis in viruses with lipid envelopes that rely on membranes for virion assembly and release. Giant viruses encode proteins involved in energy generation and nutrient acquisition across diverse habitats, suggesting that they employ conserved strategies to reprogram host metabolism (40). However, their specific contributions to host redox balance and membrane homeostasis remain poorly understood. In contrast to viruses, such as hepatitis C and dengue virus, which primarily exploit host ER enzymes to drive membrane remodeling (70, 74, 75), giant viruses encode their own metabolic enzymes−likely acquired from their hosts−to directly reprogram lipid metabolism. This distinction highlights a fundamentally different and more autonomous mechanism of host manipulation.

## Materials and methods

### Strains and growth conditions

The strains and plasmids used in this study are listed in Supplementary Table 1. Strains were grown at 37°C in NZ media or M63 minimal media supplemented with 0.2% Glucose. For DHB4 derivative strains, the minimal media was also supplemented with 50 μg/mL Leucine and 50 μg/mL Isoleucine. The antibiotic concentrations used are as following ampicillin 100 μg/mL, kanamycin 40 μg/mL, chloramphenicol 10 μg/mL, and tetracycline 10 μg/mL.

The DNA fragments from giant viruses were codon optimized for *E. coli* and synthesized using Integrated DNA Technologies (IDT) (gblocks, Supplementary Table 3). All except *Yasminevirus* sp*. vkor* sequences were fused to a 6X-His tag at the amino terminus. The plasmid backbone was amplified using PR25 with PR161 (Supplementary Table 2) using pCL113 plasmid as a template. Then, gblocks 18, and 19 were independently assembled to the vector using NEBuilder High-Fidelity assembly mix (New England Biolabs, USA) to generate plasmids PL88 and PL89. The Trc promoter of these plasmids was mutated to Trc204 using site-directed mutagenesis with primers PR242 and PR243 and the products were ligated with KLD master mix (New England Biolabs, USA) to generate PL156 and PL161. Plasmid PL161 was then used as template to amplify the vector with primers PR25 and PR161, the PCR product was then assembled with gblocks 31, 32 and 33 (Supplementary Table 3) to generate PL361, PL362, and PL363, respectively. Site directed mutagenesis to introduce two changes in the -10 box of the Trc promoter (changing it from Trc204 to Trc206) was done using primers PR176 and PR200 to generate PL357 and PL358.

To delete positive residues (R27K28K29) in Bs*vkor*, primers PR542 and PR543 were used to generate the site-directed mutagenized plasmid using plasmid PL161 as a template. Similarly, primers PR544 and PR545 together with plasmid PL156 were used to create the deletion of K26K27K28 residues in Hv*vkor*. The products were then ligated with KLD master mix (New England Biolabs, USA) to generate PL341 and PL342, respectively.

To remove cytoplasmic *dsbA* from the constructs, plasmid PL341 was amplified with primers PR90 and PR547 while PL342 was amplified with primers PR90 and PR548. The products were ligated with KLD master mix (New England Biolabs, USA) to generate PL345 and PL346, respectively.

To generate mutations that balance positive residues in Bs*vkor*, a PCR product was amplified using primers PR709 and PR710 and PL341 as template. The product was then DpnI digested and ligated using KLD master mix (New England Biolabs, USA) to generate PL453. Similarly, for Es*vkor*, a PCR product using primers PR711 and PR712 was amplified using PL361 as template and ligated as indicated above. The resulting plasmid was then used as template to amplify a second PCR product using primers PL713 and PR714 to be ligated into the final PL454. Plasmids were sequenced to verify the mutations.

### Serine auxotrophy to determine alkaline phosphatase complementation

Cytoplasmic alkaline phosphatase, when active, can substitute for the last enzyme of serine biosynthesis, serine-1-phosphate phosphatase (*serB*) (54). Thus, plasmids harboring viral VKORs were transformed into strain lacking *serB* and expressing cytoplasmic alkaline phosphatase (Δss*phoA*, DHB7787 strain). Growth was tested on M63 0.2% glucose minimal media supplemented with 50 μg/mL Leucine and 50 μg/mL Isoleucine either in the presence or absence of serine (50 μg/mL). As a positive control we used strain CL680 expressing *Aeropyrum pernix* VKORi (26).

### Motility assays

Plasmids harboring viral VKORs were transformed into two strains HK325 (76) and FSH231 (52) to test for motility complementation. Swarming assays were done in M63 0.2% glucose and 0.3% agar supplemented with antibiotics. To induce expression of viral VKORs, 10-50 µM IPTG was used in the media. As positive controls we used CL382 expressing *Mycobacterium tuberculosis* VKOR (77), and LL236 expressing human VKORc1 (53) induced with 2.5 μM and 10 μM IPTG, respectively. Bacteria were stabbed into plates and incubated at 30°C. Halos were measured after 48 h.

### Phylogenetic tree construction

Viral VKOR protein sequences were found using organism’s taxonomy (https://www.ncbi.nlm.nih.gov/Taxonomy/Browser/wwwtax.cgi). Genomic sequences for 6,200 non-redundant VKOR proteins were downloaded December 15, 2025 from UniProt (https://www.uniprot.org/) (78) using keyword:KW-1185 to select from reference genomes only. A subset of 100 protein sequences representative from each clade generated in PFAM (IPR038354 *vkor* domain superfamily, https://www.ebi.ac.uk/interpro/) was manually selected from the list (79). Five VKOR protein sequences of giant virus hosts were added to the list by searching homologs in Amoebozoa using Uniprot. A multiple sequence alignment was done with Muscle and the best substitution model was determined by maximum likelihood fit using MEGA12 (80). The phylogeny was then inferred using the Maximum Likelihood method and Whelan-Goldman WAG model (+Freq) (81) of amino acid substitutions and the tree with the highest log likelihood (-58,950.67) is shown. The percentage of replicate trees in which the associated taxa clustered together (100 replicates) is shown next to the branches (82). Initial tree(s) for the heuristic search were obtained automatically by applying the Maximum Parsimony method. The evolutionary rate differences among sites were modeled using a discrete Gamma distribution across 5 categories (+G, parameter=4.9757), with 3.48% of sites deemed evolutionarily invariant (+/). The analytical procedure encompassed 100 amino acid sequences with 1,953 positions in the final dataset. Evolutionary analyses were conducted in MEGA12 utilizing up to 4 parallel computing threads. The Newick tree generated was visualized using iTol (https://itol.embl.de/).

### Membrane topology analysis

VKOR membrane topology was determined using TOPCONS (https://topcons.cbr.su.se/) (49), TMHMM 2.0 (https://services.healthtech.dtu.dk/services/TMHMM-2.0/) and Deep TMHMM (https://services.healthtech.dtu.dk/services/DeepTMHMM-1.0/) (51). Membrane topology predictions were visualized using Protter tool (83) (https://wlab.ethz.ch/protter/start/)

### Motility suppressor analysis

Genomic DNA extraction was done using Wizard HMW extraction kit (Promega, USA) as per manufacturer instruction. DNA was then sequenced using Oxford Nanopore combined with Illumina. Sequence alignments to find single nucleotide polymorphisms, insertions, deletions were done using CLC Genomics Workbench software (Qiagen, USA).

### Random mutagenesis

To randomly mutagenize Hv*vkor,* Hv*vkor*_ΔK26K27K28_, and Ys*vkor*, a kit of Diversify™ PCR Random Mutagenesis (Takara Bio, USA) was used. The genes were amplified using PR20-PR690 or PR20-PR691 and PL156, PL342 or PL363 as templates using mutagenic conditions 1 to 5 to generate 2 to 4.6 mutations per kb. The vector was amplified by PCR with primers PR25 and PR161 using PL156 as template. A mutant pool of PCR products was made and ligated to the vector by NEBuilder High-Fidelity Assembly mix (New England Biolabs, USA). Library sizes of Hv*vkor,* Hv*vkor*_ΔK26K27K28_, and Ys*vkor* ranged between 25,000-30,000 mutants. The plasmid library was then transformed into FSH231 or DHB7787 strains. Samples of the libraries were stabbed on motility plates or plated on M63 glucose minimal media and incubated for 2 days at 30°C. Motility halos were found after incubation, while no colonies were obtained on minimal media lacking serine. Bacterial colonies were isolated from a motility halo and their plasmids were purified and transformed into FSH231 strain to confirm motility. Plasmids were then sequenced by Oxford Nanopore Sequencing to find the mutations.

### VKOR protein abundance

*E. coli* strains carrying *Harvfovirus* 6XHis-VKOR WT, Δ3 and Δ3-E95K were grown in M63 glucose minimal media with 100 µM IPTG until log phase (OD_600_ of 0.5). Proteins were precipitated with 10% TCA, washed with 600 µL of acetone and resuspended in 100 µL of Tris-HCl pH 8 with 1% SDS. Proteins were quantified with BCA protein assay (ThermoFisher Scientific). Total protein was then normalized to 20 µg, mixed with 5X-SDS reducing loading buffer, boiled for 5 min at 100°C and subjected to SDS-PAGE. Bio-Rad Mini-PROTEAN®TGX™ 12% were run for 55 min at 150 V. Proteins were then transferred onto PVDF membrane (Millipore) using Trans-Blot SD Semi Dry Transfer Cell (Bio-Rad) for 60 min at 0.08A. Membranes were then blocked with TBS (Tris Buffered Saline) containing 5% milk and incubated with 1:5,000 dilution of α-His antibody (Santa Cruz Biotechnology) overnight at 4°C. Similarly, 1:10,000 dilution of α-RpoA antibody (4RA2; BioLegend, USA) was used as a loading control. Blots were washed three times with TBS containing 0.1% Tween 20 and incubated with corresponding secondary antibody either 1:50,000 dilution of α-Mouse antibody (Santa Cruz Biotechnology) for 1 h at room temperature. Chemiluminescent substrate (ECL, Bio-Rad) was used to detect proteins using ChemiDoc™ MP Imaging System (Bio-Rad) and Amersham Hyperfilm ECL (Cytiva). Proteins were separated and transferred to membranes which were cut to blot independently with the two antibodies. The adjusted total band volume was determined for His and RpoA images using Fiji Software. The arbitrary units obtained for each band were then normalized to the volume obtained in the mutant band, which was in all cases more abundant. The relative band volume of His was then divided by the relative band volume of RpoA to obtain the relative protein abundance.

### Transcriptomic profiling of Yasminevirus and Fadolivirus infections

*Yasminevirus saudimassiliense* GU-2018 (84) and *Fadolivirus algeromassiliense* FV1/VV64 (45) strains were produced in starvation medium using *V. vermiformis* CDC19 strain as a cell host. A suspension of *V. vermiformis* was prepared by successive cycles of centrifugation (3,000 × *g* for 10 min) and resuspension in starvation medium. Triplicates of 30 mL of amoebae at 10^6^ cells/mL for each condition were inoculated with *Yasminevirus* or *Fadolivirus* at a multiplicity of infection (MOI) of 10 in a 75-cm^2^ culture flask. After 30 min of incubation at 30°C for viral entry, the supernatant was carefully aspirated to remove extracellular *Yasminevirus* or *Fadolivirus* particles and maintain amoeba’ monolayers. The cells were resuspended in 10 mL of fresh medium for a second incubation at 30°C for 16 h. This time point was considered hour zero (H0.5). At H2, H4, H6, H8, H10, H12, and H16 of each triplicate amoebae were collected and after centrifugation of 0.5 mL aliquot the sample was used for RNA extraction. A culture flask containing only amoebae in starvation medium was treated as the viral condition and was used as a negative control.

RNA extraction was performed as per manufacturer’s instructions with DNase treatement (Ozyme, Quick-RNA MiniPrep kit). mRNA was then captured with poly(A) tail using oligo(dT) coated magnetic beads following standard kit recommendations (NEBNext Poly(A) mRNA Magentic Isolation Module, New England Biolabs). After poly(A)+ RNA enrichment, a cDNA library was prepared using SuperScript IV Reverse Transcriptase (ThermoFisher), then a second brin synthesis was done using DNA Klenow I polymerase (New England Biolabs). cDNA libraries were done using COVIDSeq tests and sequenced using paired-end strategy (2*150 bp) using Illumina Novaseq Sequencer (Illumina Inc.). The Illumina integrate system was used to separate the bar-coded sequences and demultiplex paired end reads into FASTQ files.

### RNA-seq data analyses

The pre-trimmed RNAseq data was validated with FastQC (v0.11.5, (https://www.bioinformatics.babraham.ac.uk/projects/fastqc/) and MultiQC (v1.33) (85). The data was re-trimmed using fastp (v0.21.0) (86). QC and mapping metrics validated that the data was of sufficient quality and ensured the consistency of the samples. The paired-end reads were then mapped using HISAT2 (v2.2.1) (87) to the combined fasta files consisting of the *Yasminevirus saudimassiliense* GU-2018 (UPSH01000000), *Fadolivirus algeromassiliense* (MT418680.1), and the *Vermamoeba vermiformis* CDC-19 (draft genome:https://www.mediterranee-infection.com/acces-ressources/donnees-pour-articles/genome-and-annotation-of-vermamoeba-vermiformis-cdc19/) (84). The subread algorithm featureCounts was used to generate gene-level quantification (88). The data was then processed through the R (v4.5.1) package DESeq2 to identify differentially expressed genes (89). One of the triplicates of *Yasminevirus* T6 post-infection (CH6_S40) was manually excluded from the analysis because it was identified as a technical outlier. The viral gene expression patterns and heatmaps were generated in R using the pheatmap package to visualize the z-scores of the various genes across the time-ordered samples (https://github.com/raivokolde/pheatmap). Only rows with a minimum total normalized counts about 4500 and 5000, for *Yaminevirus* and *Fadolivirus*, respectively, were used. Time course comparisons between two adjacent time points were performed with R, with the later time point serving as the experimental sample and the earlier time point (or negative control) serving as the reference. The viral genes were assigned functional categories using the NCVOG dataset (https://ftp.ncbi.nih.gov/pub/wolf/COGs/NCVOG/) (90). The peptides were mapped to the NCVOG dataset using blastp (v2.6.1) with an e-value cutoff of 1×10^-5^ and with lower case masking of low complexity sequences turned on for the initial mapping.

### Proteomic profiling of Yasminevirus and Fadolivirus infections

Similar infections to the transcriptomic analysis were done for *Fadolivirus* and *Yasminevirus* cycle. One biological sample was collected at the same time points as indicated before. The pellet was subjected to enzymatic digestion into peptides using 50 μL of trypsin/LysC from the iST kit (PreOmics GmbH, #P.O.00141) on the iST APP96™ instrument, following the manufacturer’s instructions. Tryptic digestion was stopped after 3 hours of incubation. Cleaned-up samples were vacuum-dried and reconstituted in double-distilled water containing 0.1% (v/v) formic acid. Peptide concentrations were estimated by measuring absorbance at 280 nm using a NanoDrop spectrophotometer (Thermo Fisher Scientific, Wilmington, DE, USA). Peptides were analyzed by liquid chromatography–mass spectrometry (LC–MS) using a nanoElute 2 system (Bruker Daltonics) coupled online via a CaptiveSpray source to a timsTOF HT mass spectrometer (Bruker Daltonics, Bremen, Germany). A total of 1 µg of peptides was loaded onto a PepSep Xtrem C18 column (25 cm × 150 µm, 1.5 µm particle size) and separated using a 30-minute gradient. The column temperature was maintained at 50°C using a column oven (Bruker Daltonics). Peptides were ionized by electrospray using a PepSep CaptiveSpray emitter (PN: 1811107, 20 µm i.d., Bruker Daltonics) at a capillary voltage of 1600 V. Spectra were acquired in diaPASEF mode over an m/z range of 100–1700 and an ion mobility range of 0.75–1.25 Vs/cm². Protein sequences of giant viruses were retrieved from the UniProt database (release 2026_03_05) using the following search criteria: Yasminevirus (taxonomy ID: 3044860; 1,537 entries) and Fadolivirus (taxonomy ID: 2763561; 1,453 entries). DIA-PASEF data were processed using Spectronaut v20 (Biognosys) within the Proteoscape environment against viral and host libraries.

### Proteomics data analyses

The proteomic data was binned into three categories: *Yasminevirus*, *Fadolivirus*, and negative control. The limma package was used to compare both viral categories to the negative controls, determine potentially differentially abundant peptides based on a modified Student’s t-test, and visualize the results as volcano plots (91). Rows with low abundance were excluded from the analysis if the total protein abundance was less than 10, this excluded approximately 17% of the data.

### Statistical Analyses

Comparisons of protein abundance and motility halo diameter between mutants and Δ*dsbB* were done using ordinary non-paired one-way ANOVA multiple comparisons selecting for Holm-Sidáks test with GraphPad Prism version 10.6.0. Significant differences were indicated in graphs using GP style: p-value ≤0.0001 (****), 0.0002 (***), 0.021 (**), 0.0332 (*). Non-significant p-values (>0.1234) were indicated as ns.

## Author contributions

Conceptualization, D.B. and C.L.; Methodology, C.L., J.A., B.L.S.; Investigation, R.C., J.A., D.C., and C. L.; Formal Analysis, J.A., D.B.R. and C.L.; Writing – Original Draft, C.L.; Writing – Review & Editing, J.A., P.C., B.L.S. and C.L.; Visualization, D.B.R., and C.L.; Supervision and Project Administration, C.L.; Funding Acquisition, B.L.S., and C.L.

## Acknowledgements

We thank Weikai Li for helpful discussions about vitamin K carboxylases and Xindan Wang for helpful advice on genome sequence alignments. We also thank Philippe Decloquement, Saïd Azza, Vincent Bossi, Claudia Andrieu and Sofiane Bakour for generation of giant viruses transcriptomic and proteomic data.

Figures 1a, 4a, and 7a were created with Biorender.com, licenses PH29IDIP7Y, AI29IDITGS and XW29IDIM2H, respectively.

