## Supplementary Information for "Giant viruses encode vitamin K–based redox modules for lipid modification"

Running title: Vitamin K reduction coupled to lipid desaturation in giant viruses

Rebecca Collins^1^, Julien Adréani^2,3,4^, Dariana Chavez^1^, Douglas Rusch^5^, Dana Boyd^6^, Philippe Colson^2,3,4^, Bernard La Scola^2,3,4^ and Cristina Landeta^1^*

Author’s affiliations:

^1^ Department of Biology. Indiana University. Bloomington, USA.

^2^ MEPHI unit D-258, Aix-Marseille University. Marseille, France.

^3^ IHU Méditerranée Infection, Marseille, France.

^4^ Laboratoire des agents infectieux, Assistance Publique-Hôpitaux de Marseille (AP-HM), Marseille, France

^5^ Center for Genomics and Bioinformatics. Department of Biology. Indiana University. Bloomington, USA.

^6^ Harvard Medical School. Boston, USA.

**Keywords:** disulfide bond, DsbB, VKOR, VKORc1, coagulation, giant virus, epoxidase, lipid desaturation, desaturase, vitamin K carboxylase, vitamin K dependent enzyme

**Supplementary Figure 1.** Predicted topology of the membrane VKOR proteins found in giant viruses using TOPCONS consensus (https://topcons.cbr.su.se/) showing conflicted topologies.

*Barrevirus sp.* VKOR

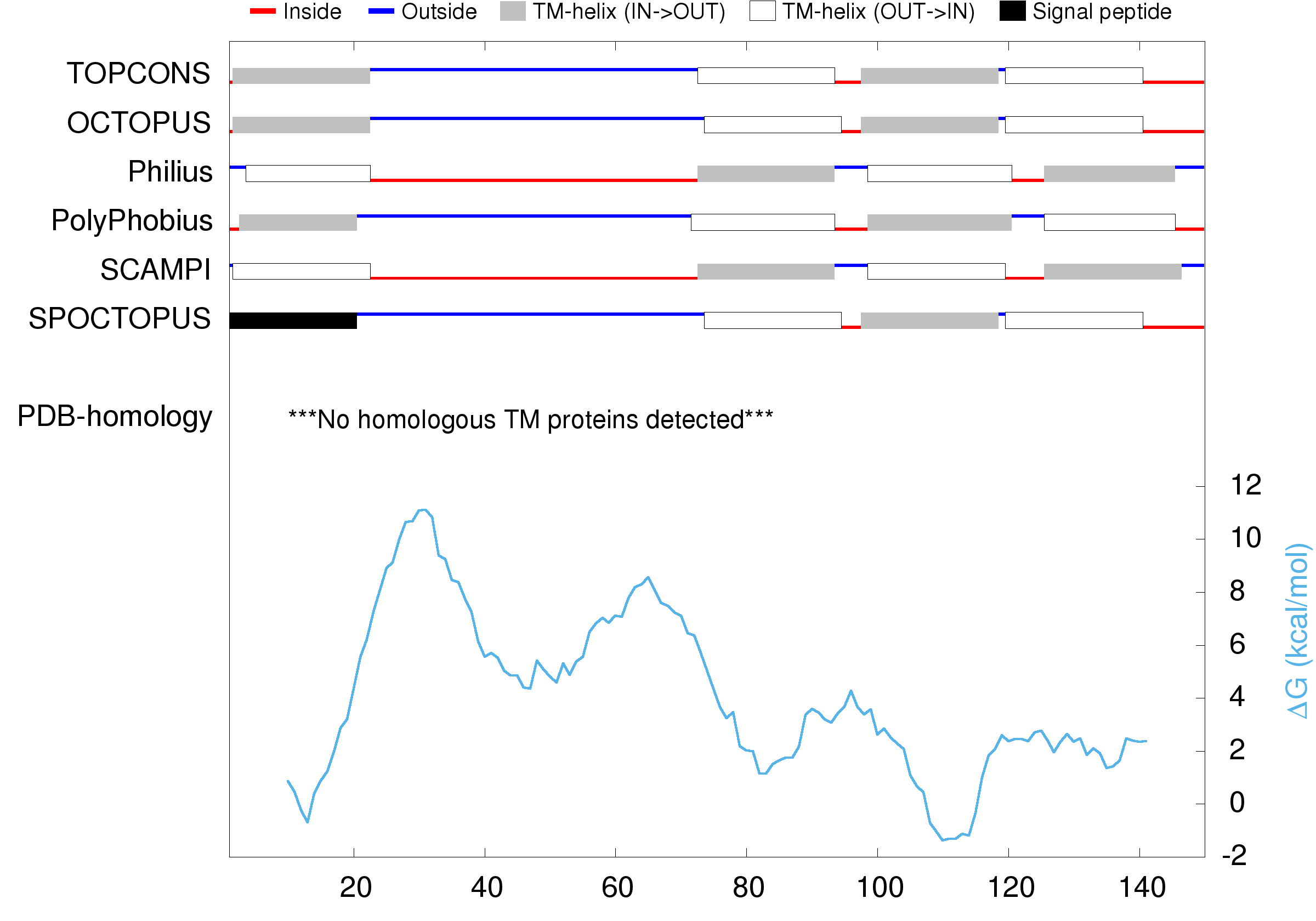

*Edafosvirus sp.* VKOR

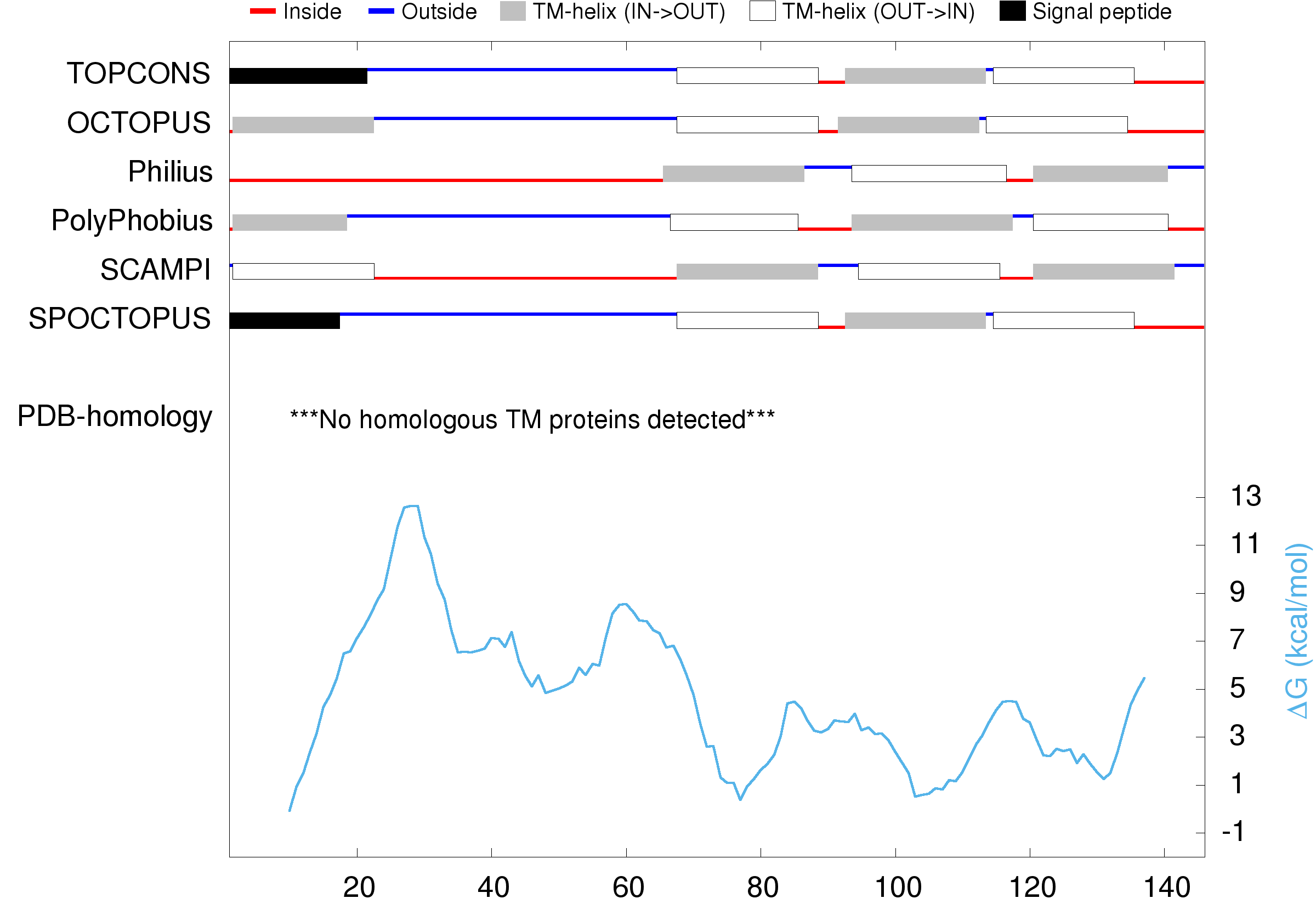

*Fadolivirus algeromassiliense* VKOR

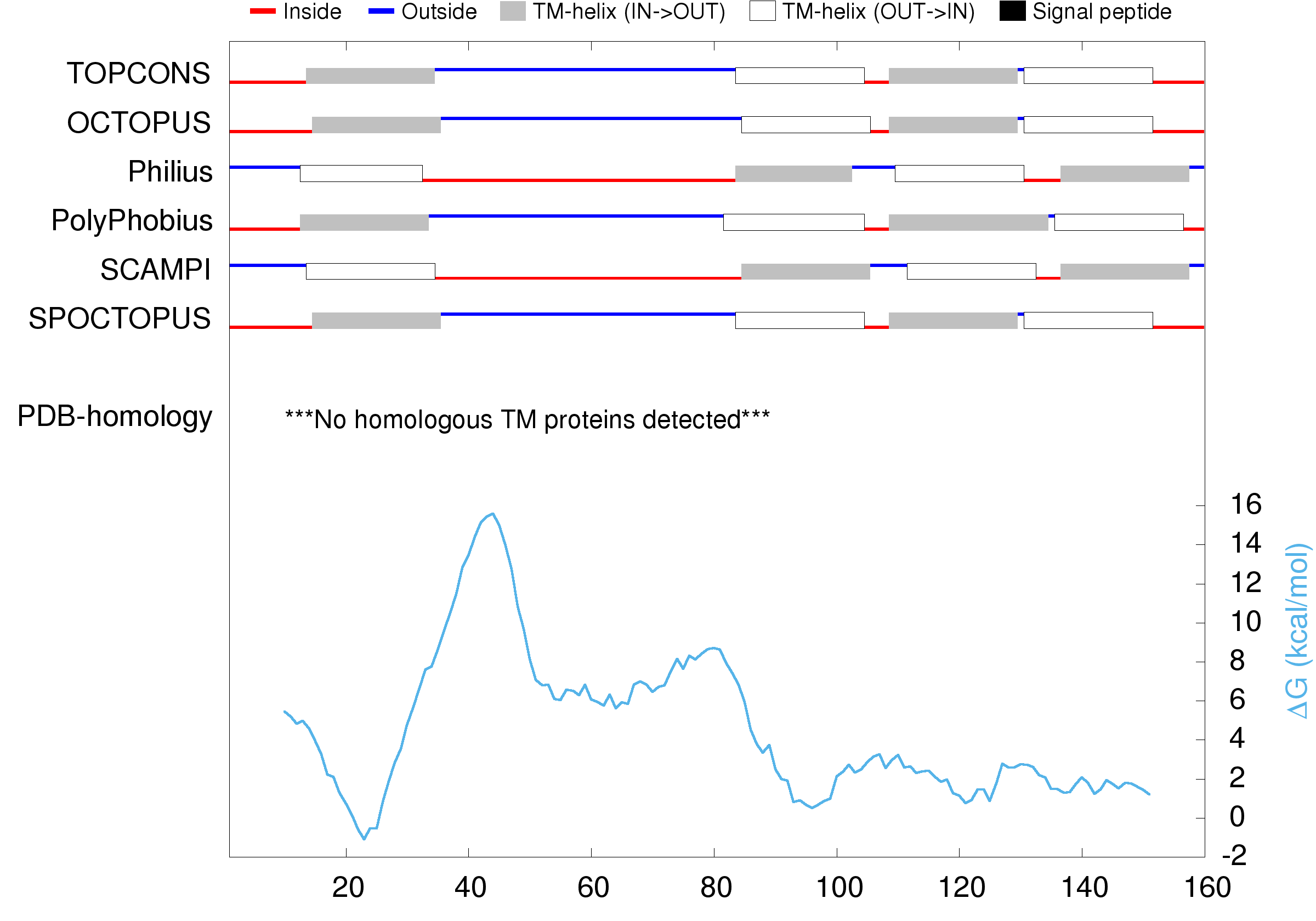

*Harvfovirus sp.* VKOR

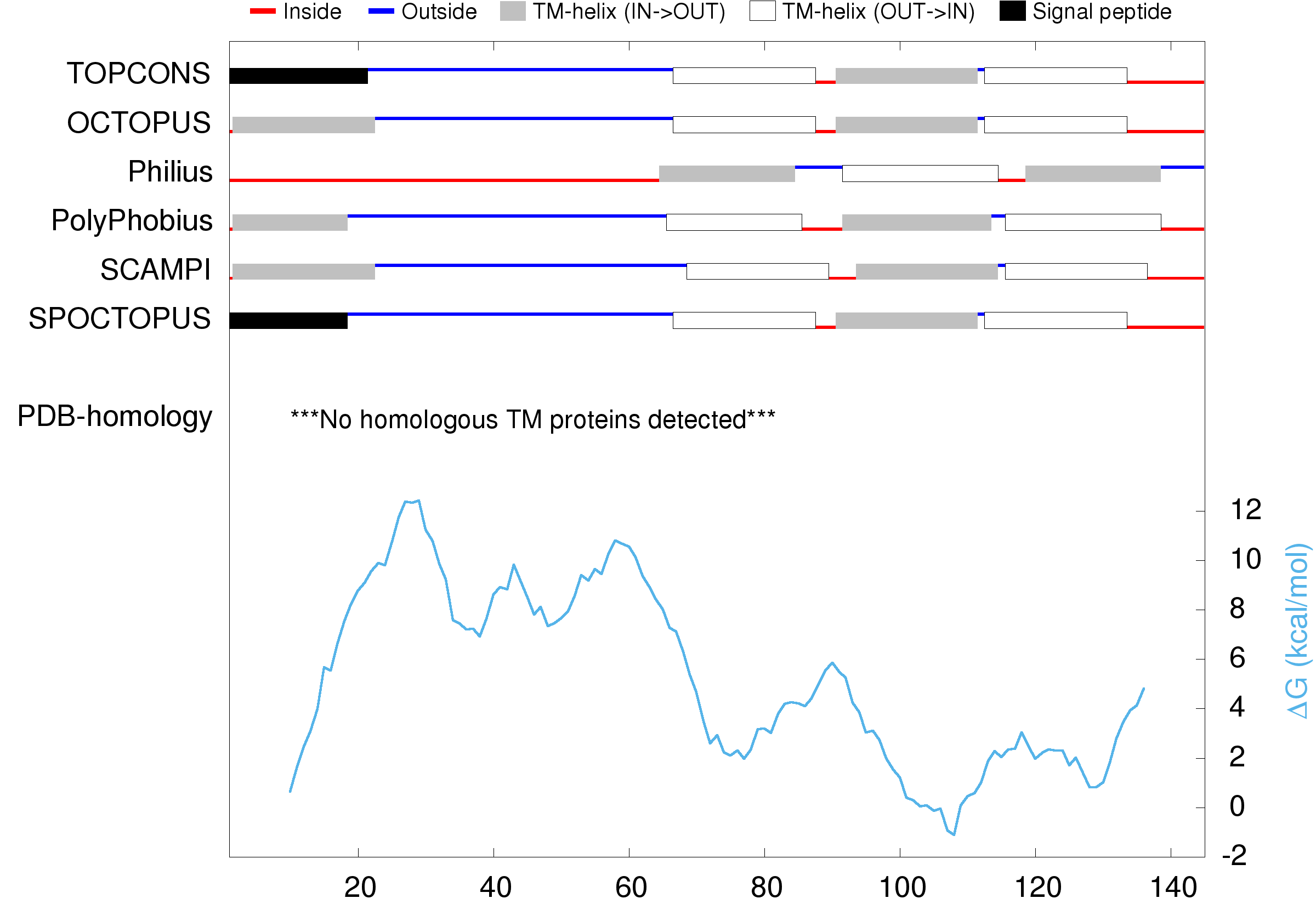

*Yasminevirus saudimassiliense* VKOR

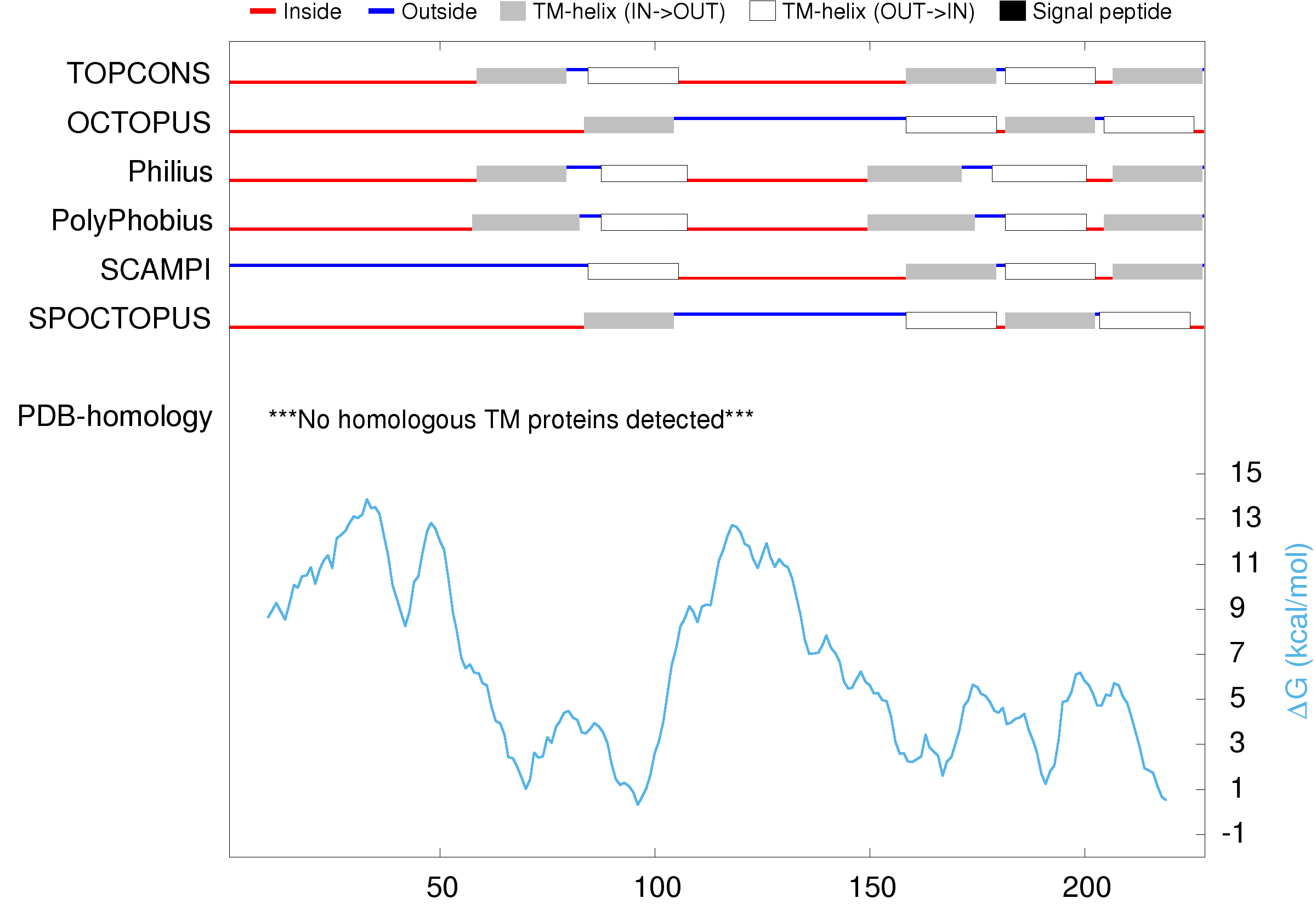

**Supplementary Figure 2.** Motility of strains with deletion of positive residues in pDSW204. Δ*dsbB** is Δ*dsbB yidC*_T362I_ *hslV*_C160Y_ alleles (FSH231 strain).

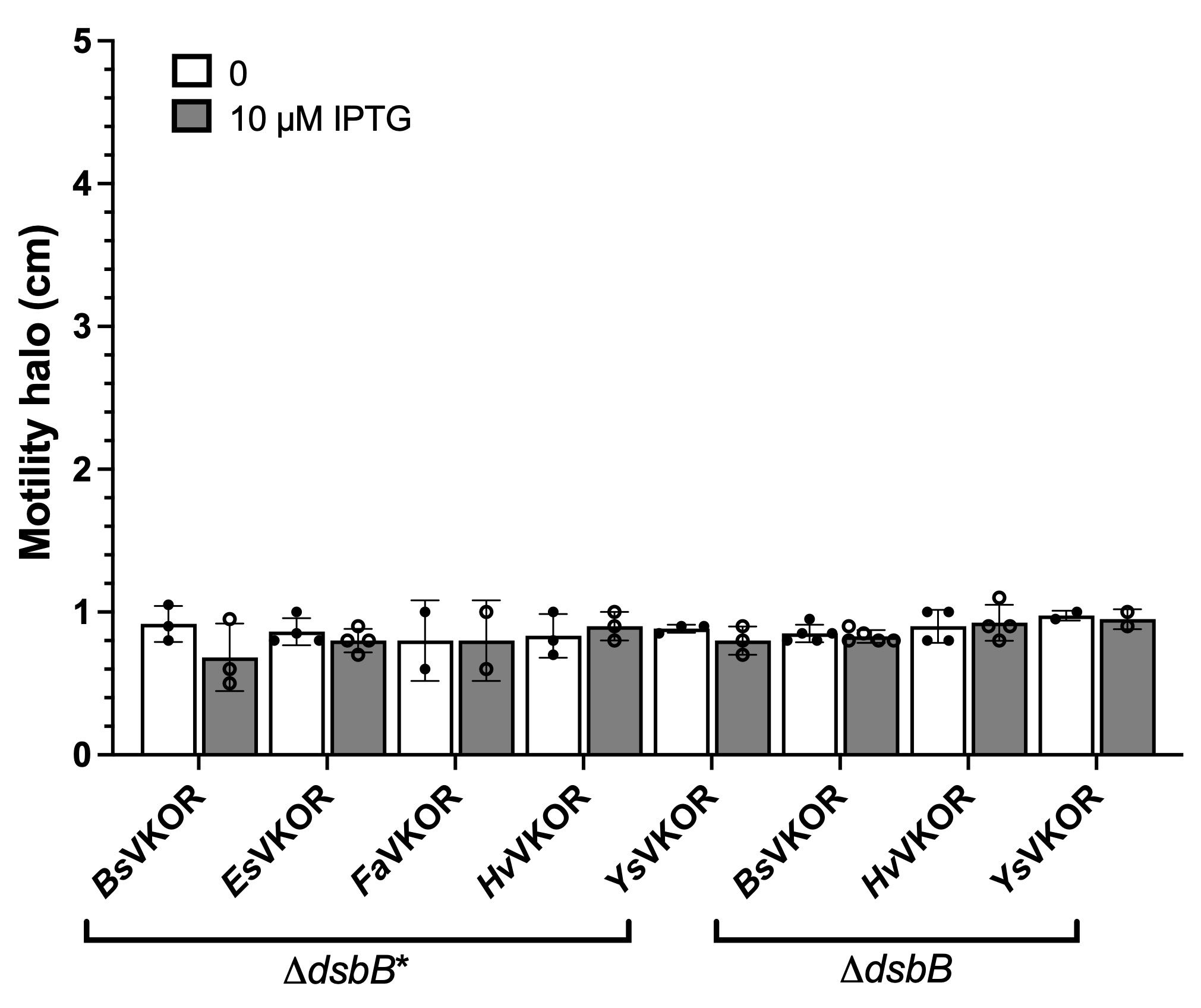

**Supplementary Figure 3.** Motility of strains with deletion of positive residues *yidC*_T362I_ *hslV*_C160Y_ alleles (Δ*dsbB**, FSH231 strain).

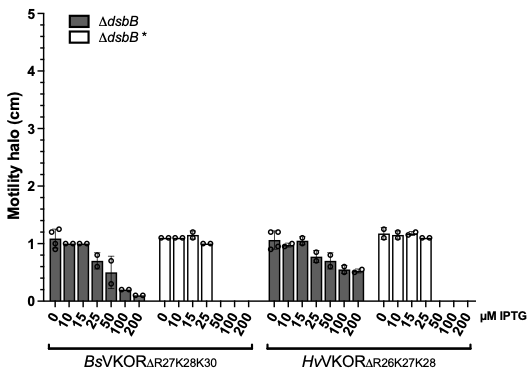

**Supplementary Figure 4.** Analysis of suppressor mutations are unrelated to viral VKOR topology. *E. coli* motility assay to test periplasmic orientation of catalytic cysteines. Suppressor mutations were isolated after 3 days of incubation with wildtype *Barrevirus*, *Harvfovirus* and *Yasminevirus* VKOR proteins. Isolated plasmids from suppressors were re-transformed in a clean background and tested for motility. Motility halos were measured after for 2 days of incubation at 30ºC. Whole genome sequencing of suppressor strains revealed OmpA loss of function mutations and a PhoQ point mutation. WT: wildtype, S: suppressor, R: re-transformed.

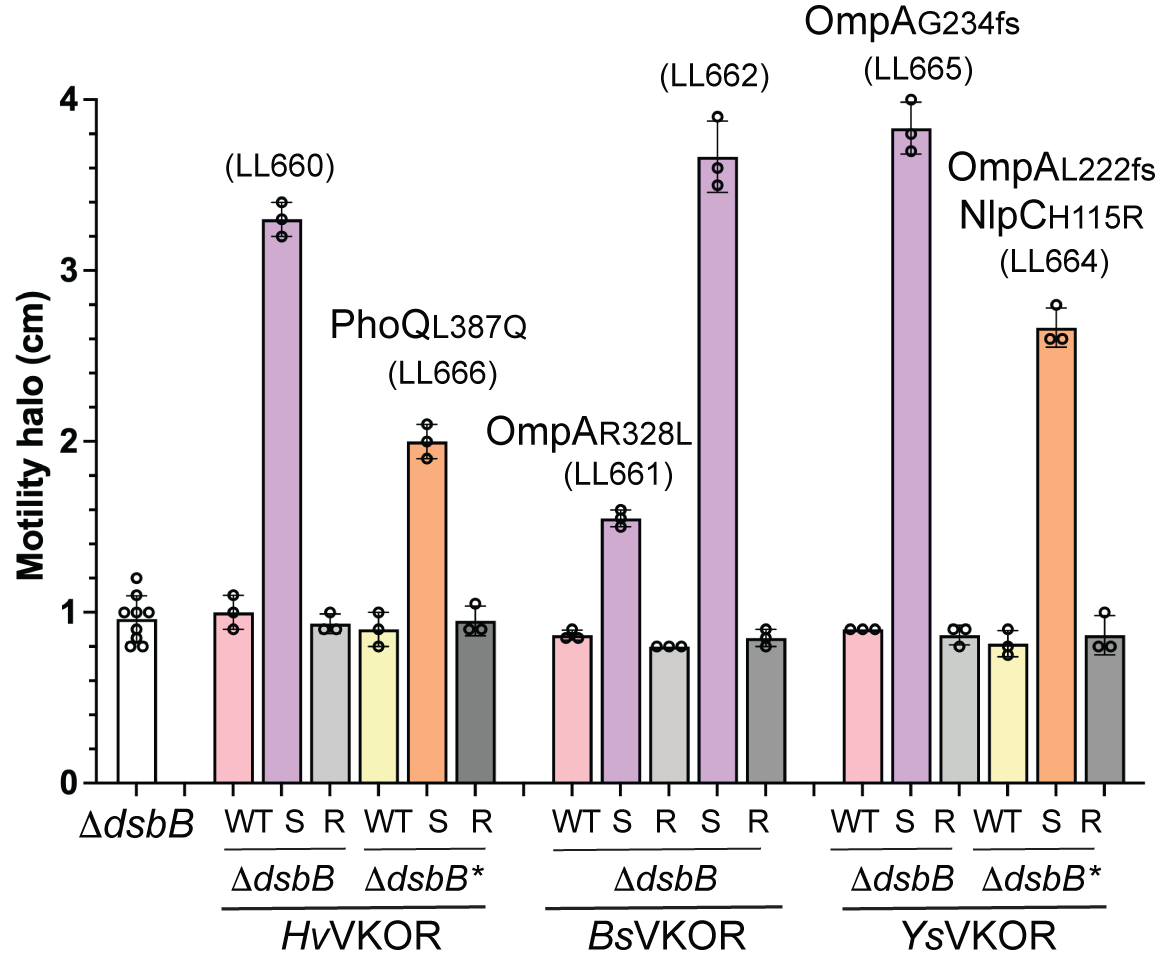

**Supplementary Figure 5.** Abundance of reads mapped against *Fadolivirus* (top), *Yasminevirus* (center), and *V. vermiformis* reference genomes. The graphs show the linear regression of three independent replicas. Principal component analysis of RNAseq dataset (bottom).

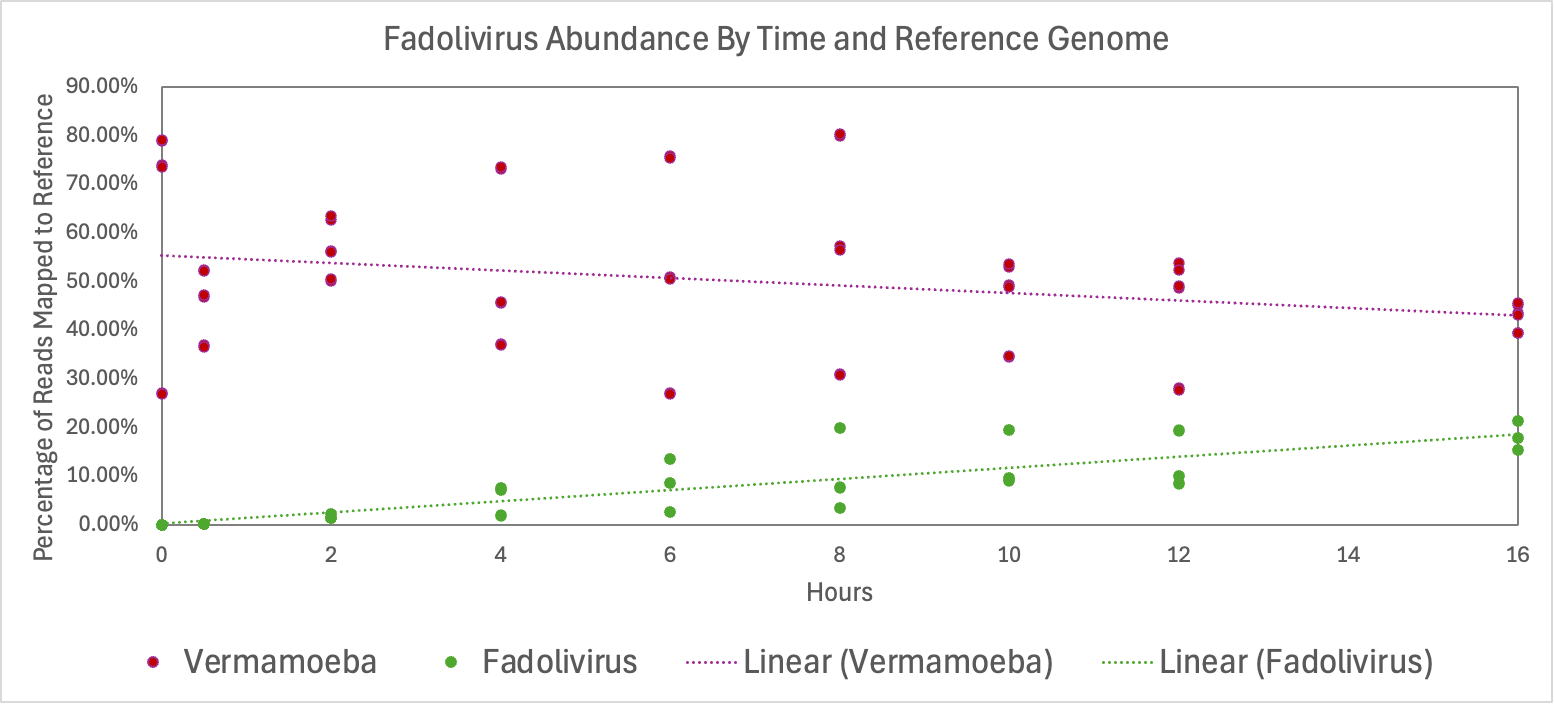

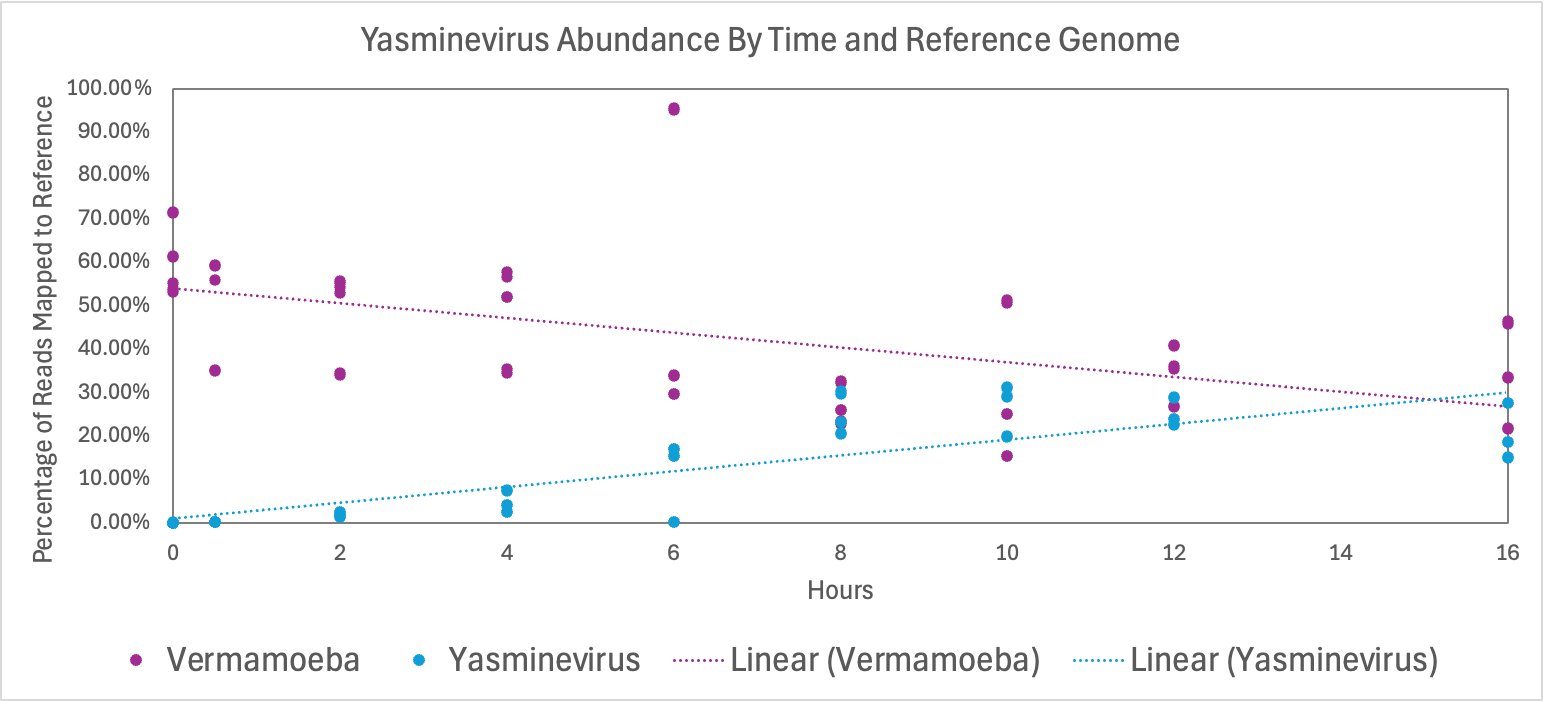

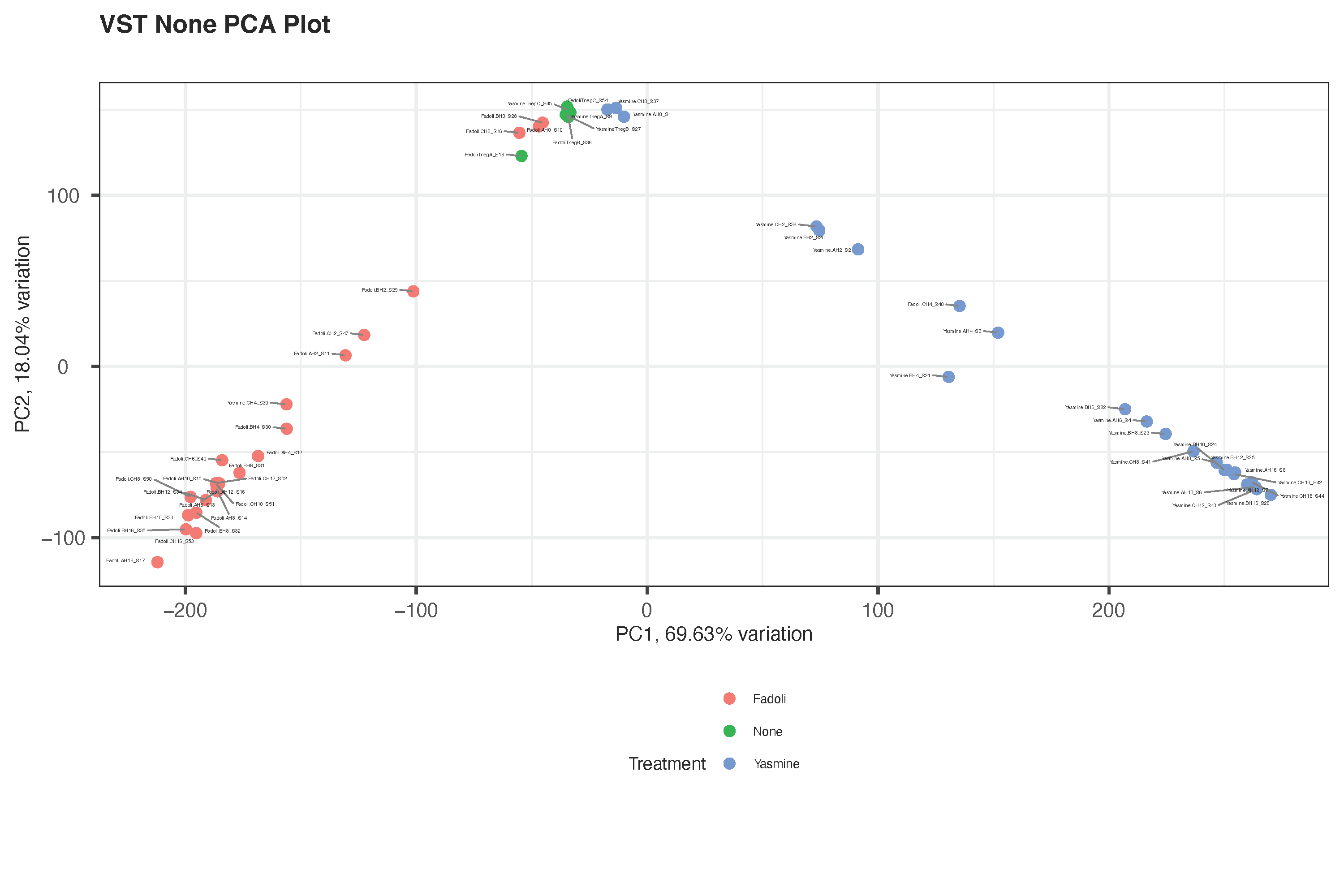

**Supplementary Figure 6**. Distribution of functional categories of *Fadolivirus* (top) and *Yasminevirus* (bottom) expressed genes when infecting *V. vermiformis*.

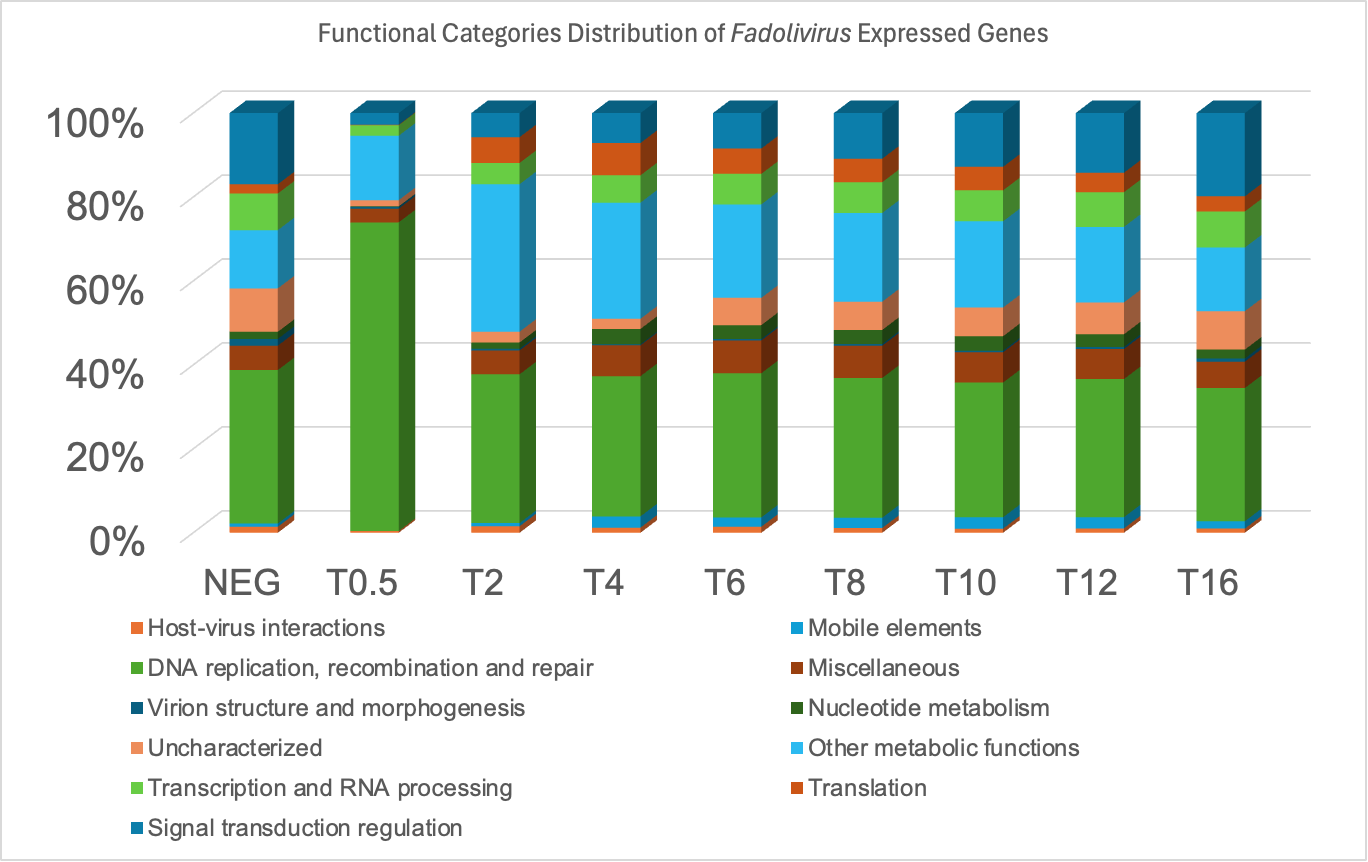

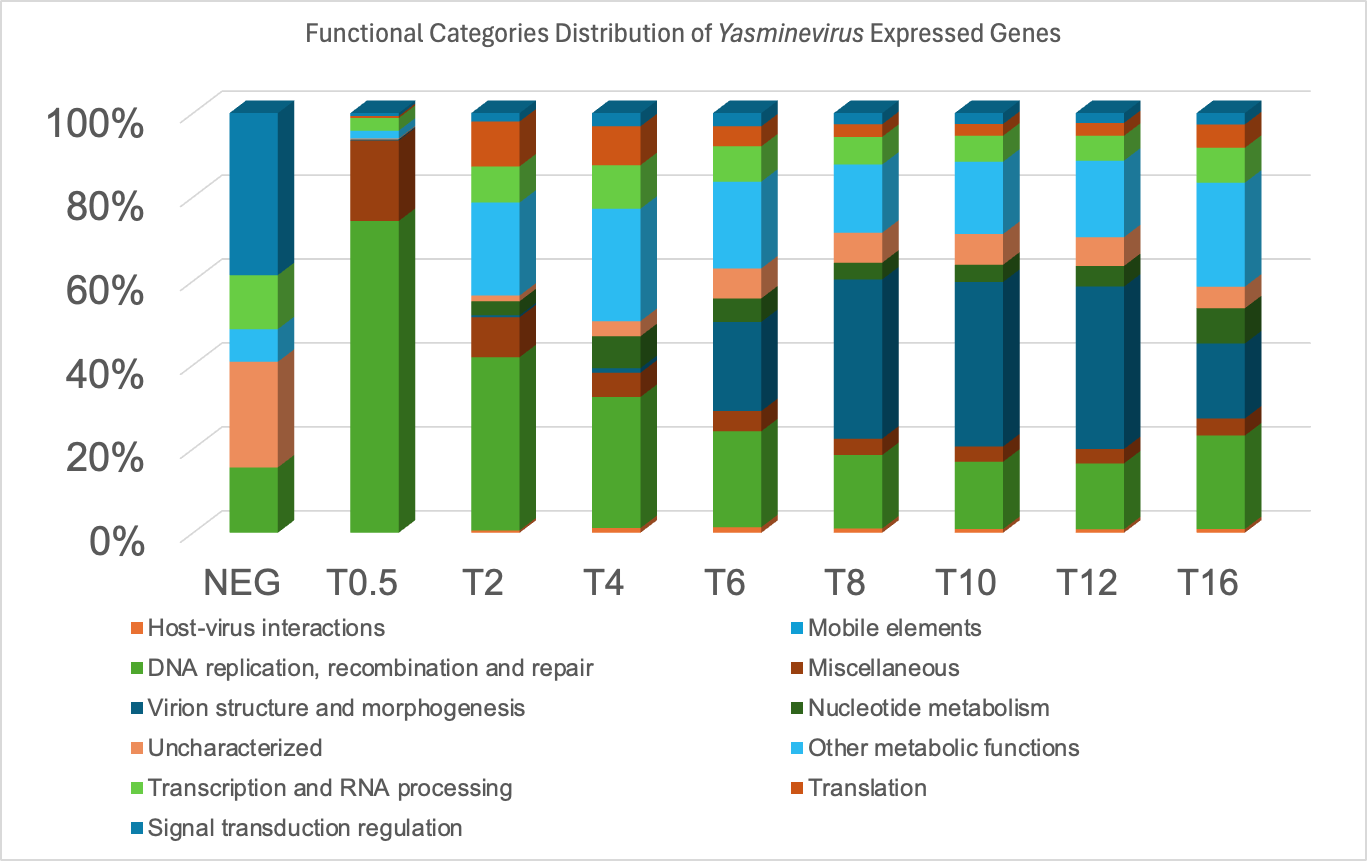

**Supplementary Figure 7**. RNAseq coverage of *Ys*VKED (*Yasminevirus*_1534) in samples of *V. vermiformis* infected with *Yasminevirus.* Three replicas of mock infected samples are shown at the top and the three replicas of *Yasminevirus*-infected at T16 are shown at the bottom.

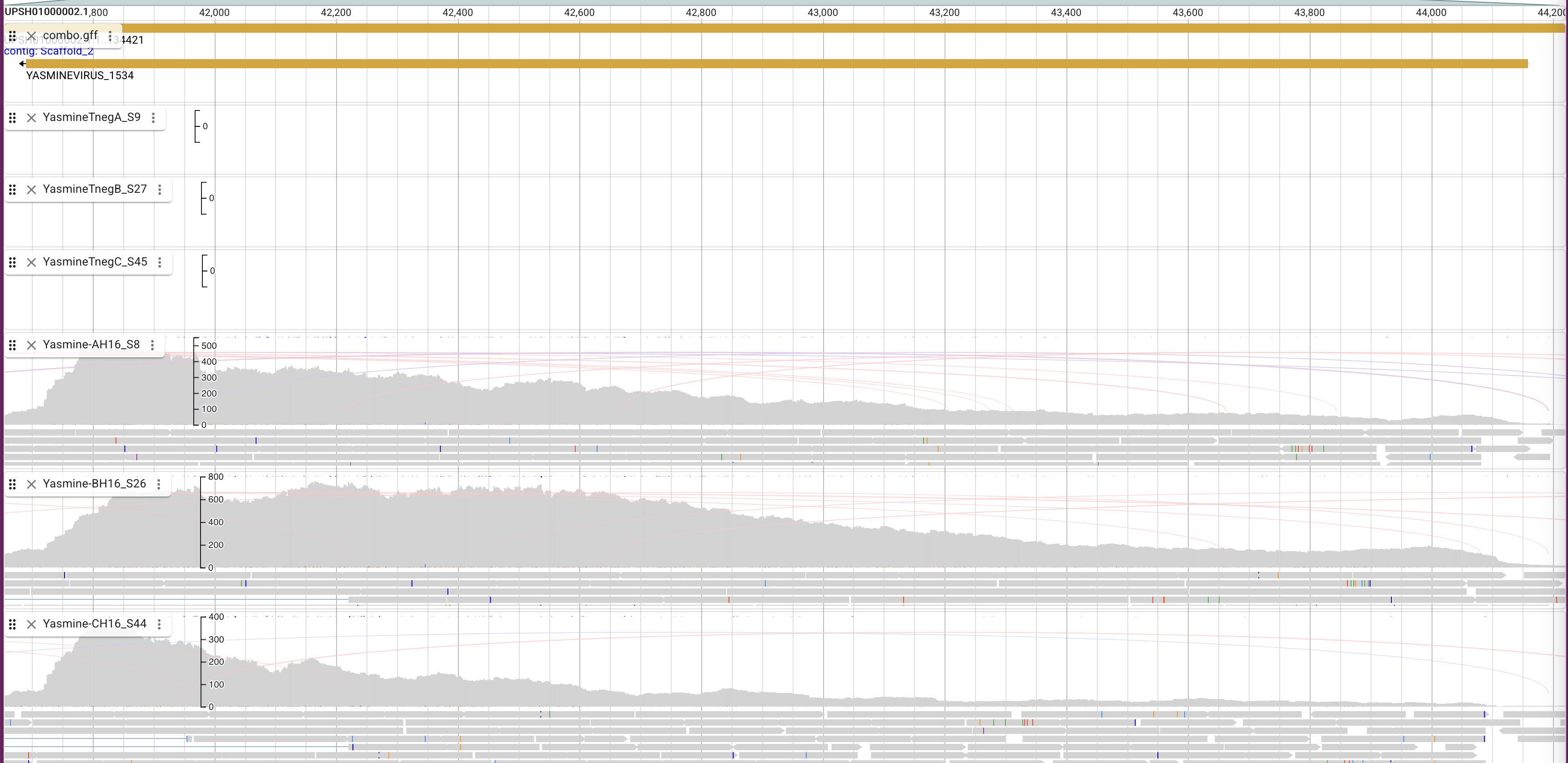

**Supplementary Tables**

**Table 1.** Strain list used in this study

| **ID** | **Genotype** | **Ref.** |
| --- | --- | --- |
| DHB4 | MC1000 Δ*phoA* PvuII *phoR* ΔmalF3 F’ [*lac*^+^ (*lacI*^Q^) *pro*] | (1) |
| HK295 | MC1000 Δ*ara*714 *leu*^+^ | (2) |
| *Yasminevirus* | *Fadolivirus* *algeromassiliense* strain FV1/VV64 | (3) |
| *Fadolivirus* | *Yasminevirus saudimassiliense* strain A1/GU-2018 | (4) |
| Amoeba | *Vermamoeba vermiformis* CDC19 | (5) |
| *E. coli* serine auxotrophs with Δss*phoA* | | |
| DHB7787 | DHB4 Δ(deoD-serB)zjj::Tn10(Tc^r^) pAD495 (Δss*phoA*, Cm^r^) | (6) |
| CL680 | DHB4 Δ(deoD-serB)zjj::Tn10(Tc^r^) pAD495 (Δss*phoA*, Cm^r^), pTrc99a-Ap*vkor_i_* -Δss*dsbA* (Amp^r^) | (6) |
| LL159 | DHB4 Δ(deoD-serB)zjj::Tn10(Tc^r^) pAD495 (Δss*phoA*, Cm^r^), pDSW204 (Amp^r^) | This study |
| LL145 | DHB4 Δ(deoD-serB)zjj::Tn10(Tc^r^) pAD495 (Δss*phoA*, Cm^r^), pTrc99a-Hv*vkor*-ΔssdsbA (Amp^r^) | This study |
| LL146 | DHB4 Δ(deoD-serB)zjj::Tn10(Tc^r^) pAD495 (Δss*phoA*, Cm^r^), pTrc99a-Bs*vkor*-ΔssdsbA (Amp^r^) | This study |
| LL213 | DHB4 Δ(deoD-serB)zjj::Tn10(Tc^r^) pAD495 (Δss*phoA*, Cm^r^), pDSW204-Hv*vkor*-ΔssdsbA (Amp^r^) | This study |
| LL214 | DHB4 Δ(deoD-serB)zjj::Tn10(Tc^r^) pAD495 (Δss*phoA*, Cm^r^), pDSW204-Bs*vkor*-Δss*dsbA* (Amp^r^) | This study |
| LL594 | DHB4 Δ(deoD-serB)zjj::Tn10(Tcr) pAD495 (ΔssphoA, Cm^r^), pDSW204-HvVKOR ΔK26K27K28 (Amp^r^) | This study |
| LL595 | DHB4 Δ(deoD-serB)zjj::Tn10(Tcr) pAD495 (ΔssphoA, Cm^r^), pDSW204-BsVKORΔR27K28K30 (Amp^r^) | This study |
| LL603 | DHB4 Δ(deoD-serB)zjj::Tn10(Tc^r^) pAD495 (Δss*phoA*, Cm^r^), pDSW206-Bs*vkor*-Δss*dsbA* (Amp^r^) | This study |
| LL604 | DHB4 Δ(deoD-serB)zjj::Tn10(Tc^r^) pAD495 (Δss*phoA*, Cm^r^), pDSW206-Bs*vkor*ΔR27K28K30-Δss*dsbA* (Amp^r^) | This study |
| LL630 | DHB4 Δ(deoD-serB)zjj::Tn10(Tc^r^) pAD495 (Δss*phoA*, Cm^r^), pDSW204-Es*vkor*-Δss*dsbA* (Amp^r^) | This study |
| LL631 | DHB4 Δ(deoD-serB)zjj::Tn10(Tc^r^) pAD495 (Δss*phoA*, Cm^r^), pDSW204-Fs*vkor*-Δss*dsbA* (Amp^r^) | This study |
| LL632 | DHB4 Δ(deoD-serB)zjj::Tn10(Tc^r^) pAD495 (Δss*phoA*, Cm^r^), pDSW204-Ys*vkor*-Δss*dsbA* (Amp^r^) | This study |
| *E. coli* strains used for motility assays | | |
| HK325 | HK295 Δ*dsbB*, λMalFlacZ102 (Km^r^) | (2) |
| FSH231 | HK295 ΔdsbB λMalFLacZ (Km^r^) YidCT362I (Cm^r^) HslVC160Y | (7) |
| CL379 | HK295 Δ*dsbB*, λMalFlacZ102 (Km^r^), pTrc99a | (8) |
| LL236 | HK295 Δ*dsbB* λMalFLacZ (Km^r^) YidCT362I (Cm^r^) HslVC160Y, pDSW204-6XHis-Hs*vkorc1*ΔA31AR N36 E37 (Amp^r^) | (9) |
| LL177 | HK295 Δ*dsbB*, λMalFlacZ102 (Km^r^), pTrc99a-Hv*vkor*-Δss*dsbA* (Amp^r^) | This study |
| LL178 | HK295 Δ*dsbB*, λMalFlacZ102 (Km^r^), pTrc99a-Bs*vkor*-Δss*dsbA* (Amp^r^) | This study |
| LL215 | HK295 Δ*dsbB*, λMalFlacZ102 (Km^r^), pDSW204-Hv*vkor*-Δss*dsbA* (Amp^r^) | This study |
| LL216 | HK295 Δ*dsbB,* λMalFlacZ102 (Km^r^), pDSW204-Bs*vkor*-Δss*dsbA* (Amp^r^) | This study |
| LL556 | HK295 Δ*dsbB,* λMalFlacZ102 (Km^r^), pDSW204-Bs*vkor*ΔR27K28K30-Δss*dsbA* (Amp^r^) | This study |
| LL557 | HK295 Δ*dsbB* λMalFlacZ102 (Km^r^), pDSW204-Hv*vkor*ΔK26K27K28- Δss*dsbA* (Amp^r^) | This study |
| LL568 | HK295 Δ*dsbB* λMalFlacZ102 (Km^r^), pDSW204-Bs*vkor*ΔR27K28K30 (Amp^r^) | This study |
| LL569 | HK295 Δ*dsbB* λMalFlacZ102 (Km^r^), pDSW204-Hv*vkor*ΔK26K27K28 (Amp^r^) | This study |
| LL572 | HK295 ΔdsbB λMalFLacZ (Km^r^) YidCT362I (Cm^r^) HslVC160Y, pDSW204-Bs*vkor*ΔR27K28K30-Δss*dsbA* (Amp^r^) | This study |
| LL573 | HK295 ΔdsbB λMalFlacZ102 (Km^r^) YidCT362I (Cm^r^) HslVC160Y, pDSW204-Hv*vkor*ΔK26K27K28-Δss*dsbA* (Amp^r^) | This study |
| LL574 | HK295 ΔdsbB λMalFLacZ (Km^r^) YidCT362I (Cm^r^) HslVC160Y, pDSW204-Bs*vkor*ΔR27K28K30 (Amp^r^) | This study |
| LL575 | HK295 ΔdsbB λMalFlacZ102 (Km^r^) YidCT362I (Cm^r^) HslVC160Y, pDSW204-Hv*vkor*ΔK26K27K28 (Amp^r^) | This study |
| LL616 | HK295 ΔdsbB λMalFlacZ102 (Km^r^) YidCT362I (Cm^r^) HslVC160Y, pDSW204-Es*vkor*-Δss*dsbA* (Amp^r^) | This study |
| LL617 | HK295 ΔdsbB λMalFlacZ102 (Km^r^) YidCT362I (Cm^r^) HslVC160Y, pDSW204-Fs*vkor*-Δss*dsbA* (Amp^r^) | This study |
| LL618 | HK295 ΔdsbB λMalFlacZ102 (Km^r^) YidCT362I (Cm^r^) HslVC160Y, pDSW204-Ys*vkor*-Δss*dsbA* (Amp^r^) | This study |
| LL619 | HK295 ΔdsbB λMalFlacZ102 (Km^r^), pDSW204-Ys*vkor*-Δss*dsbA* (Amp^r^) | This study |
| LL622 | HK295 ΔdsbB λMalFlacZ102 (Km^r^) YidCT362I (Cm^r^) HslVC160Y, pDSW204-Hv*vkor*-Δss*dsbA* (Amp^r^) | This study |
| LL623 | HK295 ΔdsbB λMalFlacZ102 (Km^r^) YidCT362I (Cm^r^) HslVC160Y, pDSW204-Bs*vkor*-Δss*dsbA* (Amp^r^) | This study |
| LL786 | HK295 ΔdsbB λMalFlacZ102 (Km^r^) YidCT362I (Cm^r^) HslVC160Y, pDSW204-6XHis-Bs*vkor*ΔR27K28K30-E95K-Δss*dsbA* (Amp^r^) | This study |
| LL788 | HK295 ΔdsbB λMalFlacZ102 (Km^r^) YidCT362I (Cm^r^) HslVC160Y, pDSW204-6XHis-Es*vkor*ΔK24K26R28-E93K-Δss*dsbA* (Amp^r^) | This study |
| *E. coli* motility suppressor strains | | |
| LL660 | HK295 Δ*dsbB*, λMalFlacZ102 (Km^r^), pDSW204-Hv*vkor*-Δss*dsbA* (Amp^r^) | This study |
| LL661 | HK295 Δ*dsbB,* λMalFlacZ102 (Km^r^), pDSW204-Bs*vkor*-Δss*dsbA* (Amp^r^) | This study |
| LL662 | HK295 Δ*dsbB,* λMalFlacZ102 (Km^r^), pDSW204-Bs*vkor*-Δss*dsbA* (Amp^r^) | This study |
| LL663 | HK295 ΔdsbB λMalFlacZ102 (Km^r^) YidCT362I (Cm^r^) HslVC160Y, pDSW204-Es*vkor*-Δss*dsbA* (Amp^r^) | This study |
| LL664 | HK295 ΔdsbB λMalFlacZ102 (Km^r^) YidCT362I (Cm^r^) HslVC160Y, pDSW204-Ys*vkor*-Δss*dsbA* (Amp^r^) | This study |
| LL665 | HK295 Δ*dsbB,* λMalFlacZ102 (Km^r^), pDSW204-Ys*vkor*-Δss*dsbA* (Amp^r^) | This study |
| LL666 | HK295 ΔdsbB λMalFlacZ102 (Km^r^) YidCT362I (Cm^r^) HslVC160Y, pDSW204-Hv*vkor*-Δss*dsbA* (Amp^r^) | This study |
| LL761 | HK295 ΔdsbB λMalFlacZ102 (Km^r^) YidCT362I (Cm^r^) HslVC160Y, pDSW204-Hv*vkor*ΔK26K27K28-E91K-Δss*dsbA* (Amp^r^) | This study |
| Plasmids | | |
| pTrc99a | Trc promoter, pBR322 origin (Amp^r^) |  |
| pDSW204 | Promoter down mutation in -35 of pTrc99a (Amp^r^) | (10) |
| pDSW206 | Promoter down mutation in -10 of pTrc99a (Amp^r^) | (10) |
| pAD495 | pACYC ori, pTac-*phoA*Δ2-22 (Δss*phoA*, Cm^r^) | Derman A. |
| pCL113 | pTrc99a-Ap*vkor_i_* -*dsbA*Δ1-19 (Δss*dsbA,* Amp^r^) | (6, 10) |
| PL88 | pTrc99a-6XHis-Hv*vkor*-Δss*dsbA* (Amp^r^) | This study |
| PL89 | pTrc99a-6XHis-Bs*vkor*-Δss*dsbA* (Amp^r^) | This study |
| PL156 | pDSW204-6XHis-Hv*vkor*-Δss*dsbA* (Amp^r^) | This study |
| PL161 | pDSW204-6XHis-Bs*vkor*-Δss*dsbA* (Amp^r^) | This study |
| PL341 | pDSW204-6XHis-Bs*vkor*ΔR27K28K30-Δss*dsbA* (Amp^r^) | This study |
| PL342 | pDSW204-6XHis-Hv*vkor*ΔK26K27K28-Δss*dsbA* (Amp^r^) | This study |
| PL345 | pDSW204-6XHis-Bs*vkor*ΔR27K28K30 (Amp^r^) | This study |
| PL346 | pDSW204-6XHis-Hv*vkor*ΔK26K27K28 (Amp^r^) | This study |
| PL357 | pDSW206-6XHis-Bs*vkor*-Δss*dsbA* (Amp^r^) | This study |
| PL358 | pDSW206-6XHis Bs*vkor*ΔR27K28K30-Δss*dsbA* (Amp^r^) | This study |
| PL361 | pDSW204-6XHis-Es*vkor*-Δss*dsbA* (Amp^r^) | This study |
| PL362 | pDSW204-6XHis-Fs*vkor*-Δss*dsbA* (Amp^r^) | This study |
| PL363 | pDSW204-Ys*vkor*-Δss*dsbA* (Amp^r^) | This study |
| PL453 | pDSW204-6XHis-Bs*vkor*ΔR27K28K30-E95K-Δss*dsbA* (Amp^r^) | This study |
| PL454 | pDSW204-6XHis-Es*vkor*ΔK24K26R28-E93K-Δss*dsbA* (Amp^r^) | This study |

**Table 2**. Primers used in this study

| **ID** | **Primer sequence** |
| --- | --- |
| PR20 | GAGCGGATAACAATTTCACACAGG |
| PR25 | ggtctgtttcctgtgtgaaattgttatccgctcacaattc |
| PR161 | ggtacccggggatccAGGAGATATA |
| PR176 | CATCCGGCTCGCATTATGTGTGGAATTGTGAGCG |
| PR200 | ATTAATTGTAAACAGCTCATTTCAGAATATTTGCCAG |
| PR242 | ATGAGCTGTTTACAATTAATCATCC |
| PR243 | TTCAGAATATTTGCCAGAAC |
| PR542 | GGCACTACGCTCGACGTAGATGGCG |
| PR543 | AGCCGCTATGTCGCTGTGTGCGATATG |
| PR544 | TTTTTGTGCGACATCAATGAAAATA |
| PR545 | CAGGTGGGCGTTCTTCTCTACATAA |
| PR90 | ggctgttttggcggatgag |
| PR547 | TCAGTAGAACAGTTCACTGAATGCC |
| PR548 | TCAACTAATCAGACTATTTTCTTTCAAT |
| PR690 | TCCCCGGGTACCTCAACTAATCAGACTATTTTCTTTC |
| PR691 | TCCCCGGGTACCTCAATATAAGAAGTAGATCAGTGTTAAA |
| PR709 | CCGCAAGGTCCTGTTTTTCGGAGCT |
| PR710 | AAAGGGATAATCGTAAAGGGATAGATAGG |
| PR711 | TTAAGAAGTTCCTGCTTCTGGTCGCAAGTT |
| PR712 | ATGGAATCAAAGTAAAAGGGTACATCTTG |
| PR713 | GAGGTTAACGCTCACCAGGCCCTTTGCGACTTGAACGA |
| PR714 | AACATAGATGGCATATGCCGACACAAGAACGCC |

**Table 3**. Codon-optimized DNA fragments encoding viral VKOR proteins. A 6X-His tag followed by a thrombin site (underlined) was added at the amino terminus for all except *Ys*VKOR.

| **ID** | **Organism** | **Sequence** |
| --- | --- | --- |
| *Hv*VKOR (g19) | *Harvfovirus* sp. AYV81228.1 | gaaacagaccATGGGCAGCAGCCATCATCATCATCATCACAGCAGCGGCCTGGTGCCGCGCGGCAGCCATATGATCTCCCTGTTGGGTTTGGCCGGGATCATCATTAGTTCTTACGCTGTTTATGTAGAGAAGAACGCCCACCTGAAGAAGAAATTTTTGTGCGACATCAATGAAAATATGTCCTGCTCACGTATTCTGACAAGCGATTACAGCAAGATGGTTGAGATGATCTTCAAGCTGAAACCGAAGCACCCTCTGAACTTGCCCAATACTTATTATGGAATTCTGTTTTATATCATTGTCACGTTGTACCCTCATTTCCAGATGATTCCCTTTCGTGAATACCTTTTTTTCACGGGCTCCTTACTGTCTATGCTTGCCTCTATCACACTTATCTGCATTTTAATCTTTAAGTTAAAAGAATTGTGCATTGTATGCTTAGCTACACACTTGATCAATATGTGCATTTTTTATTTGGCATTGAAAGAAAATAGTCTGATTAGTTGAggtacccggg  MGSSHHHHHHSSGLVPRGSHMISLLGLAGIIISSYAVYVEKNAHLKKKFLCDINENMSCSRILTSDYSKMVEMIFKLKPKHPLNLPNTYYGILFYIIVTLYPHFQMIPFREYLFFTGSLLSMLASITLICILIFKLKELCIVCLATHLINMCIFYLALKENSLIS* |
| *Bs*VKOR (g18) | *Barrevirus* sp*.* AYV77270.1 | gaaacagaccATGGGCAGCAGCCATCATCATCATCATCACAGCAGCGGCCTGGTGCCGCGCGGCAGCCATATGAGTTTGATTTCAGCCGCCTTGTCCTATATGGACTCAACGTCGTTCGTGGTCTCATTATGCTGTATCGGGATTTTGGTGTGCTGGTACGCAATGGACATTGAGAGCGGGGAAAAGAAACCGAAGTACAAGCGTATGTGTGACGTTAACGACTCGATGTCCTGTACCCTTGTACTGACATCAAAGTATGGTCATATGGCTAAATTGATGTTCGGCTTGGATCGCAACTCACTTTTTAACCGTAGCAACGCAGAGTATGGGTTCGTTTTCTATCTTGGCCTGCTTATTTTTCAGTTCTACCCCTTCACGATGTTACCATTCTACAATTATATCTTCCTTATCGGCACGGTCGGGTCAGTATGCGCGTCCATCGGCCTGGCGTGGATTCTTTATAGCATTCTGCATAACTTCTGCATGATTTGTGTATGCATGTATGCGGTGAACACCTTATTGATGATCTCTGCAATCATGCGCGTAATGTGAggtacccggg  MGSSHHHHHHSSGLVPRGSHMDNIISLLGLIGIAISLYAIYVERSARKSKRYVAVCDMNESVSCSLVLISAYSKLGEVYLGLSKDSIFNLPNSYYGILFYIAITIYPIYPFTIIPFREVLFFGASILSIGVCCLLAWILYFKLNNFCAICATTYILNMFILYQAFSELFY* |
| *Es*VKOR  (g31) | *Edafosvirus* sp*.* AYV77849.1 | gaaacagaccATGGGCAGCAGCCATCATCATCATCATCACAGCAGCGGCCTGGTGCCGCGCGGCAGCCATATGATTACCTTGCTGGGGATTTGTGGCGTTCTTGTGTCGGCATATGCCATCTATGTTGAGGTTAACGCTAAACACAAGCAGCGTGCCCTTTGCGACTTGAACGAAGGAATGTCTTGCACGCGCGTTCTTACATCACCCTATGCTCGCATGACGGGGTTAGTATTCGGGCTGAAGAAGTCGCATCCATTAAACTTACCCAACACCTATTACGGTCTTCTGTTTTACGTGGCCATCATCCTGTACAAGATGTACCCTTTTACTTTGATTCCATTTAAGGAGTTCCTGCTTCTGGTCGCAAGTTCAATGAGTATGGCGGCATGTATTGGTCTTGCATATGTGCTTTACTTCAAATTGAAGGATATTTGTATCGTCTGCATCACAACCTATATCATTAATTCGTGCATCTTCTATTATGCCCTTAAAGAAAACAACATTATTTGAggtacccggg  MGSSHHHHHHSSGLVPRGSHMITLLGICGVLVSAYAIYVEVNAKHKQRALCDLNEGMSCTRVLTSPYARMTGLVFGLKKSHPLNLPNTYYGLLFYVAIILYKMYPFTLIPFKEFLLLVASSMSMAACIGLAYVLYFKLKDICIVCITTYIINSCIFYYALKENNII  MITLLGICGVLVSAYAIYVEVNAKHKQRALCDLNEGMSCTRVLTSPYARMTGLVFGLKKSHPLNLPNTYYGLLFYVAIILYKMYPFTLIPFKEFLLLVASSMSMAACIGLAYVLYFKLKDICIVCITTYIINSCIFYYALKENNII |
| *Fa*VKOR (g32) | *Fadolivirus* *algeromassiliense* QKF94487.1 | gaaacagaccATGGGCAGCAGCCATCATCATCATCATCACAGCAGCGGCCTGGTGCCGCGCGGCAGCCATATGAGTTTGATTTCAGCCGCCTTGTCCTATATGGACTCAACGTCGTTCGTGGTCTCATTATGCTGTATCGGGATTTTGGTGTGCTGGTACGCAATGGACATTGAGAGCGGGGAAAAGAAACCGAAGTACAAGCGTATGTGTGACGTTAACGACTCGATGTCCTGTACCCTTGTACTGACATCAAAGTATGGTCATATGGCTAAATTGATGTTCGGCTTGGATCGCAACTCACTTTTTAACCGTAGCAACGCAGAGTATGGGTTCGTTTTCTATCTTGGCCTGCTTATTTTTCAGTTCTACCCCTTCACGATGTTACCATTCTACAATTATATCTTCCTTATCGGCACGGTCGGGTCAGTATGCGCGTCCATCGGCCTGGCGTGGATTCTTTATAGCATTCTGCATAACTTCTGCATGATTTGTGTATGCATGTATGCGGTGAACACCTTATTGATGATCTCTGCAATCATGCGCGTAATGTGAggtacccggg  MGSSHHHHHHSSGLVPRGSHMSLISAALSYMDSTSFVVSLCCIGILVCWYAMDIESGEKKPKYKRMCDVNDSMSCTLVLTSKYGHMAKLMFGLDRNSLFNRSNAEYGFVFYLGLLIFQFYPFTMLPFYNYIFLIGTVGSVCASIGLAWILYSILHNFCMICVCMYAVNTLLMISAIMRVM |
| *Ys*VKOR (g33) | *Yasminevirus saudimassiliense* GU-2018 VBB19004.1 | gaaacagaccATGAGTAGCCAGAGCCTGGATATGTCCGCAGAGATCACAACTGAGCTGCGTCAACGCAAAAGTTTTCTTCCTGTAACACAAGATGAAAAAGAACAGAAACTGCAATTTGACGCGTCCTCAGCGGAATCTCAATCGTGTGCGCGTCGTAAACGCGAATCGACCGCGAGCCTTTTCATGAAAAGTTTGAAGTTCTTTTCGTTCGTATTCGTCGTCGCCGGCGTAGCCTCAGTCGCGATGGAACACTTCTTGGGCTGCTCGCGCATTTACCTTTACTTCCTGATCGGGTTATATGTGTCTTCCCTGGCCTCCTATTATAAGTATCGCGTATGGATGGACCCTAGCTATCGTCCCGACTGTGACTGCGCCCAACCCGAGCCCGATACGTTCATCCCGACTACGCAGAACATGATGGACGGAGTTTTTACTGTTTTAGCACATAAGAAGTCAGCGTTGTTGTTAGGGATCCCAAACACAGTTTTCGGCATGGTCTTCTACACTGGTATGATTTTGATCAACTATTATAAGTTTCCGTTCTGCCACGAATTGACACTTTTATCAACCATTGTCTCATGCACAGGCAGCGTTTATCTGTGGTACACAATGGTCAATGAGGTTCGCTCAGTTTGCGTGATTTGTAGTAGTATTCACGCGATCTCATTTTTAACACTGATCTACTTCTTATATggtacccggg  MSSQSLDMSAEITTELRQRKSFLPVTQDEKEQKLQFDASSAESQSCARRKRESTASLFMKSLKFFSFVFVVAGVASVAMEHFLGCSRIYLYFLIGLYVSSLASYYKYRVWMDPSYRPDCDCAQPEPDTFIPTTQNMMDGVFTVLAHKKSALLLGIPNTVFGMVFYTGMILINYYKFPFCHELTLLSTIVSCTGSVYLWYTMVNEVRSVCVICSSIHAISFLTLIYFLY |

Supplementary Tables 4-6 are provided in excel.

**Supplementary Table 4.** RNA sequence mapping, comparison to negative across all times and gene clustering

**Supplementary Table 5.** Protein abundances of V. vermiformis samples infected with *Fadolivirus* (0.5-16h) and mock infected control (negative)

**Supplementary Table 6.** Protein abundances of *V. vermiformis* samples infected with *Yasminevirus* (0.5-16h) and mock infected control (negative)

**Supplementary Table 7.** Peptide ion intensities of *V. vermiformis* samples infected with *Fadolivirus* (0.5-16h) and mock infected control (negative)

**Supplementary Table 8.** Peptide ion intensities of *V. vermiformis* samples infected with *Yasminevirus* (0.5-16h) and mock infected control (negative)

**Supplementary Table 9.** Protein differential abundance analysis of *V. vermiformis* samples infected with *Yasminevirus* and *Fadolivirus* compared to the mock infected control (negative)
